# Single-cell analyses identify circulating anti-tumor CD8 T cells and markers for their enrichment

**DOI:** 10.1101/2020.09.30.294959

**Authors:** Kristen E. Pauken, Osmaan Shahid, Kaitlyn A. Lagattuta, Kelly M. Mahuron, Jacob M. Luber, Margaret M. Lowe, Linglin Huang, Conor Delaney, Jaclyn M. Long, Megan E. Fung, Kathleen Newcomer, Katy K. Tsai, Melissa Chow, Samantha Guinn, Juhi R. Kuchroo, Kelly P. Burke, Jason M. Schenkel, Michael D. Rosenblum, Adil I. Daud, Arlene H. Sharpe, Meromit Singer

## Abstract

The ability to monitor anti-tumor CD8^+^ T cell responses in the blood has tremendous therapeutic potential. Here, we used paired single-cell RNA sequencing and T cell receptor (TCR) sequencing to detect and characterize “tumor matching” (TM) CD8^+^ T cells in the blood of mice with MC38 tumors and melanoma patients using the TCR as a molecular barcode. TM cells showed increased activation compared to non-matching T cells in blood, and appeared less exhausted than matching counterparts in tumor. Importantly, PD-1, which has been used to identify putative circulating anti-tumor CD8^+^ T cells, showed poor sensitivity for identifying TM cells. By leveraging the transcriptome we identified candidate cell surface marker panels for TM cells in mice and melanoma patients, and validated NKG2D, CD39, and CX3CR1 in mice. These data demonstrate that the TCR can be used to identify tumor-relevant populations for comprehensive characterization, reveal unique transcriptional properties of TM cells, and develop marker panels for tracking and analysis of these cells.

**Summary:** Using single-cell RNA-sequencing coupled with TCR sequencing, we detected CD8^+^ T cell clones shared between blood and tumor in mice and melanoma patients, characterized these matching clones in blood and tumor, and identified potential biomarkers for their isolation in blood.

## Introduction

Cancer immunotherapy has revolutionized treatment of many solid and liquid tumors, delivering durable responses in some patients, as well as decreased toxicity in many instances compared to conventional cytotoxic therapies (Chen and Mellman, 2017; Ribas and Wolchok, 2018; Sharma and Allison, 2015; Sharpe and Pauken, 2017; Sun et al., 2018). CD8^+^ T cells are a central component of protective anti-tumor immune responses, and many immunotherapies are aimed at boosting the anti-tumor activity of these cells. Both preclinical and clinical data have highlighted the importance of the systemic immune response to anti-tumor immunity following checkpoint blockade (Fransen et al., 2018; Huang et al., 2019; Huang et al., 2017; Kamphorst et al., 2017; Sharpe and Pauken, 2017; Spitzer et al., 2017; Valpione et al., 2020; Wei et al., 2019; Wu et al., 2020). Recent evidence in cancer patients suggests that recruitment of new CD8^+^ T cells from the circulation into the tumor, a concept termed “clonal replacement,” may be associated with better responses to immunotherapy (Cloughesy et al., 2019; Valpione et al., 2020; Wu et al., 2020; Yost et al., 2019). The vasculature is a major site of CD8^+^ T cell trafficking between secondary lymphoid organs, primary tumors, and metastatic sites (Masopust and Schenkel, 2013; Mueller et al., 2013), making the blood an ideal location to sample to interrogate peripheral anti-tumor responses. A number of elegant studies have provided comprehensive profiling of T cells in the blood of cancer patients, including during checkpoint blockade (Chalabi et al., 2020; Huang et al., 2019; Huang et al., 2017; Kamphorst et al., 2017; Twitty et al., 2020; Valpione et al., 2020; Wei et al., 2019; Wei et al., 2017; Wu et al., 2020). However, improved methods to identify T cell populations directed against tumor epitopes are needed to focus these analyses to the minority of the circulating T cell population that has prognostic and functional relevance to the anti-tumor immune response.

Tracking antigen-specific T cell populations in the blood is difficult for a number of reasons, including the small number of T cells specific for each antigen and limited reagents to track these cells (Jenkins et al., 2010; Martinez and Evavold, 2015). Tetramers have been the gold standard for identifying antigen-specific T cells, but have a number of limitations, including: (1) the antigen must be known, (2) there are a limited number of Major Histocompatibility Complex (MHC) haplotypes with existing tetramer reagents, and (3) tetramers can be inefficient at binding low affinity T cell receptors (TCRs) (Jenkins et al., 2010; Martinez and Evavold, 2015). In humans, select surrogate markers including Programed Death 1 (PD-1, also known as CD279) and B-and-T Lymphocyte Attenuator (BTLA) have been used to try to capture the anti-tumor response in blood (Gros et al., 2016; Gros et al., 2014; Huang et al., 2019; Huang et al., 2017; Kamphorst et al., 2017; Twitty et al., 2020; Yan et al., 2018). PD-1 is used because it is associated with chronic antigen stimulation and T cell exhaustion observed in cancer; however, PD-1 is at least transiently expressed on all T cells following activation, and is not an exhaustion-specific marker (Sharpe and Pauken, 2017; Wherry and Kurachi, 2015). Moreover, a population of PD-1^+^ T cells can be found in the blood of healthy people (Duraiswamy et al., 2011). Additionally, use of such markers makes an assumption about the differentiation state of the population of interest, and consequently has the potential to introduce biases in the populations being tracked. Overcoming these obstacles to allow routine, unbiased tracking of tumor-specific T cell populations in the blood would bring substantial statistical power and biological precision to analyses of tumor-relevant T cell populations, and provide much needed insight to the overarching immune response to cancer.

Here we asked whether single cell RNA sequencing of the TCR could be used to track tumor-relevant T cell responses in the blood. Using the TCR as a “molecular barcode”, we utilized paired tumor and blood samples to identify tumor-matching (TM) blood CD8^+^ T cells that had shared TCRs with the tumor infiltrating lymphocyte (TIL) population in mice with MC38 tumors and melanoma patients. TM cells showed increased signs of activation compared to non-TM cells in the blood, but were less dysfunctional than corresponding clones in the tumor. Interestingly, in longitudinal samples from two patients that failed to respond to checkpoint blockade, the TM peripheral blood CD8^+^ T component dramatically shifted to present a stronger dysfunctional signature than before. By leveraging the transcriptional profile of TM cells, we identified a number of candidate surface markers that enrich for the TM component in the blood, and validated three of these candidate markers at the protein level using CITE-seq technology (Stoeckius et al., 2017) in mice. Importantly, combinations of these marker genes achieved improved performance compared to single markers at identifying TM cells. This work presents a general approach to deeply characterize peripheral blood T cell populations that are associated with an ongoing immune response to tumor, and to identify marker panels to enable focused and statistically powered analyses of such populations of interest.

## Results

### Identification of an activated population of CD8^+^ T cells in the blood with TCRs that match to CD8^+^ T cells in MC38 tumors

Considering the clinical relevance of monitoring anti-tumor CD8^+^ T cell responses in the blood, we investigated ways in which tumor antigen-specific T cells could be tracked in the blood of tumor-bearing mice. We first assessed PD-1 protein expression on CD8^+^ T cells in mice that had received a subcutaneous challenge with highly immunogenic colon adenocarcinoma MC38 cells. PD-1 levels were uniformly high on CD8^+^ T cells in tumors of these mice; however, PD-1 levels on CD8^+^ T cells in the blood were extremely low (Fig. 1A), casting doubt on the comprehensiveness by which PD-1 expression might be capturing the tumor-relevant CD8^+^ T cell component.

**Figure 1:**
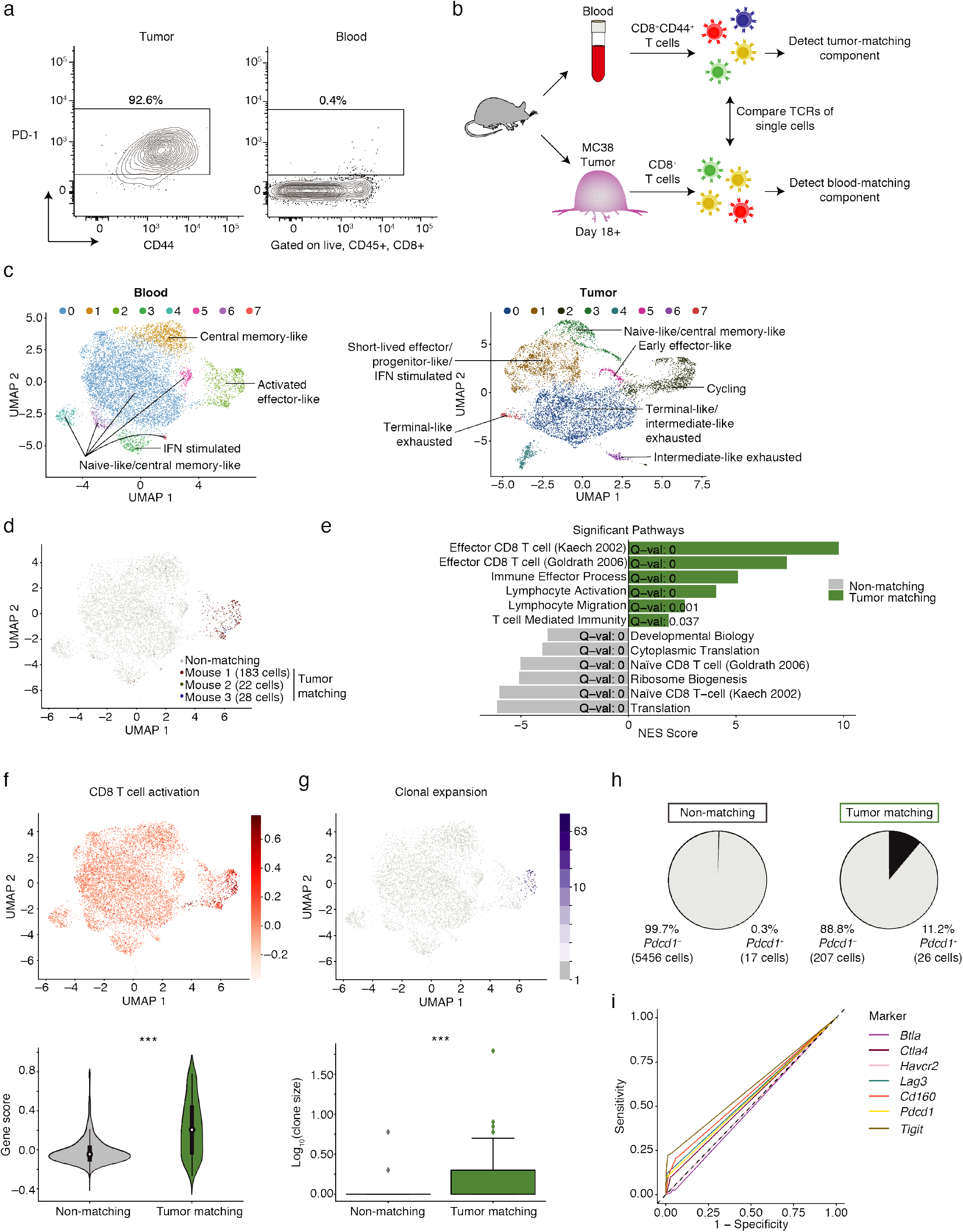
Use of single-cell RNA seq of CD8^+^ T cells identifies MC38 tumor-matching clones in blood based on TCR sequence. (a) Flow cytometry contour plots showing PD-1 and CD44 protein expression on CD8^+^ T cells in MC38 tumors and paired peripheral blood on day 21 post tumor cell implantation. Gated on singlets, live, CD45^+^, CD8α^+^ cells. Frequency of the parent population expressing PD-1 protein shown. Data are representative of 4 independent experiments with n=5-9 mice per experiment. (b) Schematic showing experimental design for single cell RNA seq. (c) Clustering and UMAP visualization of paired blood (n=10,289 cells) and MC38 tumors (n=8,450 cells) on day 18^+^ post tumor cell implantation. Data integrated from three mice from two independent experiments (exp. 1 = M1, exp. 2 = M2, M3). Colors indicate transcriptional clusters. Labels marking each cluster indicate the phenotypic description of the cluster based on gene expression data (see Methods). Up regulated genes in each cluster are in Table S1. (d) UMAP showing CD8^+^ T cells that have a TCR matching to CD8^+^ T cells found in tumor (referred to as TM cells), colored by each mouse. Grey indicates CD8^+^ T cells that do not have a TCR matching to T cells in tumor (referred to as non-TM). Cells were classified as matching if both the alpha and beta chain were identical. (e) Selected gene signatures significantly associated with genes up regulated in TM cells or non-TM cells in the integrated blood sample. Significance determined using a GSEA PreRanked analysis (Subramanian et al., 2005). Full list is in Table S3. (f) UMAP showing expression of a CD8^+^ T cell activation signature across cells in the integrated blood sample (Top) (Sarkar et al., 2007). Violin plots quantifying this enrichment in the TM cells compared to non-TM cells (Bottom). Significance determined using a Wilcoxon rank-sum test, p=1.1×10^−41^. (g) UMAP showing clonal expansion based on the number of cells with a given TCR for the integrated peripheral blood sample (Top). Box plot quantifying clonal expansion in TM cells compared to non-TM cells shown above (Bottom). Boxes show the first quartile, median, and third quartile, while the whiskers cover 1.5 times the interquartile range. Significance determined using a Wilcoxon rank-sum test, p=4.6×10^−7^. (h) Frequency of *Pdcd1*^+^ cells in the integrated blood sample separated by TM and non-TM cells. (i) ROC curve showing the sensitivity and specificity of *Pdcd1, Btla, Ctla4, Havcr2, Lag3, Cd160,* or *Tigit* to distinguish TM cells from non-TM cells. The calculated AUCs are as follows: *Pdcd1*=0.548, *Btla*=0.486, *Ctla4*=0.535, *Havcr2*=0.500, *Lag3*=0.556, *Cd160*=0.574, *Tigit*=0.603. The dashed line (on the diagonal) represents the sensitivity and specificity values of random chance.

Since the TCR encodes the specificity of a T cell for its antigen, we hypothesized that the TCR sequence could be used to assess which clones in blood were relevant to the anti-tumor response. To address this question, we performed single-cell RNA sequencing with paired TCR sequencing on CD8^+^ T cells isolated from the peripheral blood and MC38 tumors from three mice between two independent experiments (Fig. 1B, S1A-F). We reasoned that the sensitivity provided by this approach would provide a less biased method to identify blood CD8^+^ T cells that shared TCRs with TIL, referred to as “tumor matching” or TM cells. Because we expected the TM cells in the blood to be rare, we sorted for CD44^+^ CD8^+^ T cells to enrich for antigen-experienced populations in the blood. Although all CD8^+^ T cells sorted from blood expressed some level of CD44, the cells from mouse 1 (experiment 1) were sorted on CD44^high^, while the cells from mouse 2 and mouse 3 (experiment 2) included both CD44^mid^ and CD44^high^ cells. Samples were computationally integrated and the single cell transcriptomic landscapes of sorted CD44^+^ CD8^+^ T cells in blood (n=10,289 cells) and bulk CD8^+^ T cells in tumors (n=8,540 cells) were characterized (Fig. 1C). Phenotypes were defined based on up regulated genes for each cluster (Table S1) and expression of key markers (see Methods) (Fig. S1E, S1F). In the blood, the majority of the cells recovered had a naïve-like and/or central-memory like phenotype (Fig. 1C, Table S1) as expected in the CD44^mid-high^ population of specific pathogen-free (SPF) mice (Beura et al., 2016). Additional phenotypes identified in the blood corresponded with recent interferon (IFN) stimulation and an activated effector-like subpopulation (Fig. 1C, Table S1). In the tumor, more diversity was observed in the differentiation states detected, including a number of exhausted subpopulations consistent with the recently described progenitor and terminal subpopulations (He et al., 2016; Im et al., 2016; Kurtulus et al., 2019; Miller et al., 2019; Sade-Feldman et al., 2018; Siddiqui et al., 2019; van der Leun et al., 2020), as well as an intermediate-like subpopulation (Fig. 1C, Table S1). Additionally, clusters were identified in the tumor that were consistent with naïve and/or central memory-like, effector-like, cycling, and IFN-stimulated states (Fig. 1C, Table S1) (Best et al., 2013; Kakaradov et al., 2017; Milner et al., 2017). These data highlight the diversity of CD8^+^ T cell differentiation states possible in MC38 tumors, particularly in comparison to the differentiation states present in blood (Fig. 1C, Table S1).

To assess clonal overlap between CD8^+^ T cells in blood and tumor, we (a) exclusively compared T cells with at least one alpha and one beta chain, and excluded cells that did not meet this criterion (Fig. S1G), and (b) classified cells to be from the same clone if they exactly matched in their TCR sequences. Using the TCR as a molecular barcode, we observed a population of blood CD8^+^ TM cells that shared TCRs with CD8^+^ T cells in the tumor (Fig. 1D, S1A, S1D). These TM cells were transcriptionally unique from CD8^+^ T cells in blood that did not have TCRs matching to clones in the tumor (referred to as non-TM cells). Differential gene (DE) expression analysis showed elevated expression of a number of activation markers (including *Ccl5, Gzmb, Gzma, Klrg1, Itgal, Klrk1,* and *Cx3cr1*) and decreased expression of naïve-like and/or central-memory like markers (including *Ccr7, Sell, Lef1, Il7r,* and *Tcf7*) in TM cells compared to non-TM cells (Table S2). A pathway enrichment analysis of these genes showed an enrichment in TM cells for effector CD8^+^ T cell signatures, immune effector processes, and lymphocyte migration, while non-TM cells showed enrichment for naïve CD8^+^ T cell signatures (Fig. 1E, Table S3). Additionally, an analysis of manually curated transcriptional signatures from the literature showed that TM cells are enriched for activation and tissue resident memory (TRM) signatures, while non-TM cells were enriched for quiescent (naïve and/or central memory-like) signatures (Fig. 1F, S1I-S1J, Table S3). TM cells were also more likely to be clonally expanded than non-matching CD8^+^ T cells in the blood (Fig. 1G), though signatures of cell cycle were low in the blood (Fig. S1H). Importantly, the *Pdcd1* transcript provided poor sensitivity for recovering TM cells. Here, 11.2% of TM cells expressed the *Pdcd1* transcript (Fig. 1H), suggesting that if PD-1 were used as a sole marker to identify TM cells in blood, then a significant fraction of TM cells would fall below the limit of detection. Using receiver operating characteristic (ROC) curves, *Pdcd1* as well as other inhibitory receptors (including *Havcr2, Lag3, Cd160, Tigit,* and *Btla*) performed poorly in distinguishing TM cells from non-TM cells, nearing the level of random chance (Fig. 1I). Collectively, these data are consistent with blood TM CD8^+^ T cells showing an active immune response against the tumor, and support the use of the TCR as a molecular barcode to identify this population of interest rather than relying solely on individual markers such as PD-1.

### The transcriptional signature of tumor-matching CD8^+^ T cells in the blood can be used to identify markers for enrichment via flow cytometry

Given our observation that TM CD8^+^ T cells had a distinct transcriptional profile compared to non-TM cells in blood, we hypothesized that a machine learning classifier could be trained to predict if a given CD8^+^ T cell from blood is TM or non-TM based on transcriptional data. We observed that a regularized logistic regression classifier could successfully predict if a cell was TM or non-TM based on the transcriptional data for a cell, achieving high sensitivity and specificity (Fig. 2A, cross validated AUC=0.99). We next asked whether a classifier using information from only the cell surface genes would be sufficient to distinguish TM from non-TM cells, to assess the potential of identifying cell surface marker panels for flow cytometry-based sorting for downstream applications. We observed that classifiers utilizing only a list of cell-surface genes (Chihara et al., 2018) also achieved high sensitivity and specificity (Fig. 2B, cross validated AUC=0.985).

**Figure 2:**
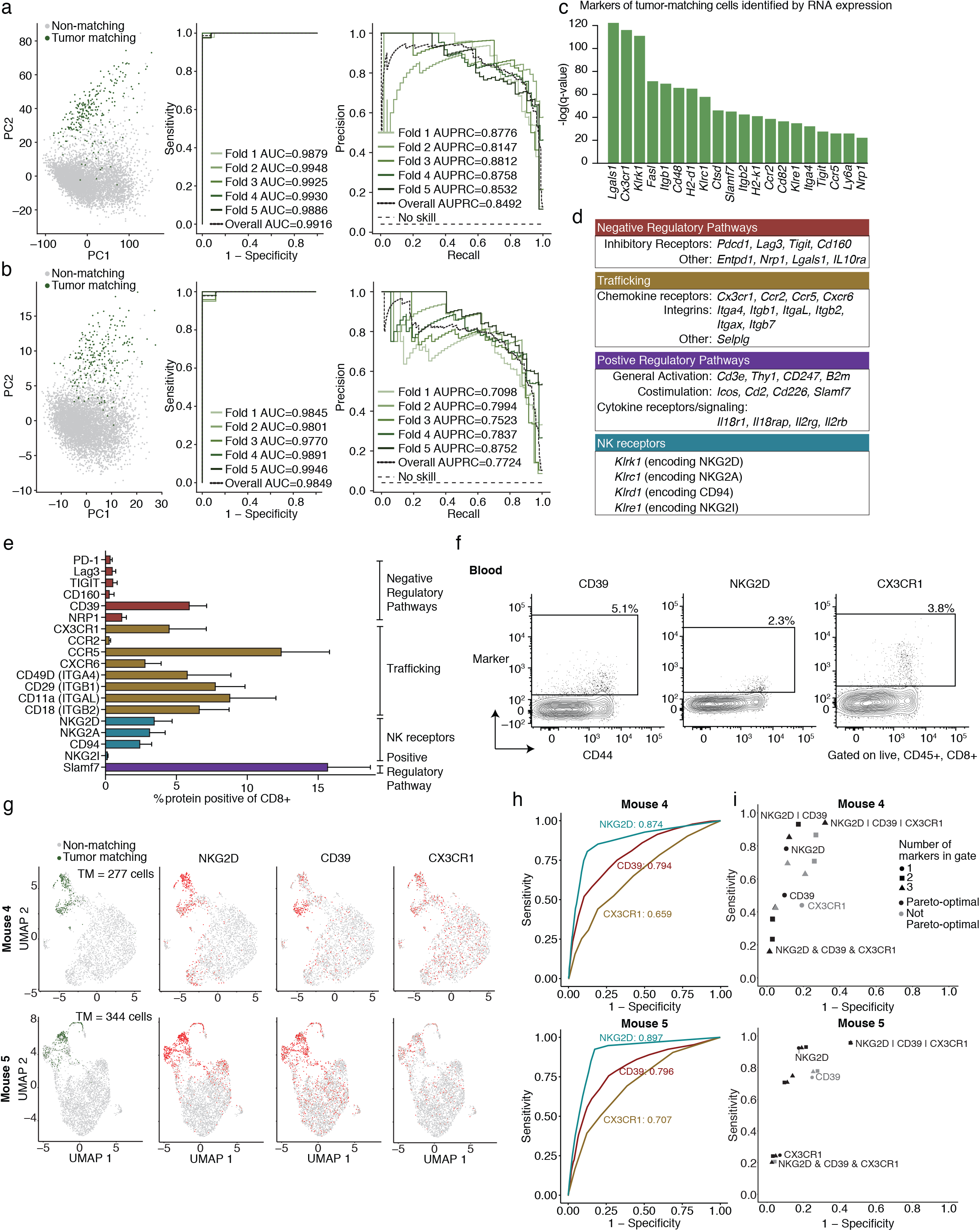
Cell-surface marker panels can be used to enrich tumor-matching cells from blood. (a-b) Logistic regression showing classification of cells as TM or non-TM based on (a) all genes and (b) a pre-selected list enriched for surface-marker genes (Chihara et al., 2018). Shown are the first two principal component projections (left), ROC curves (middle), and the Recall-Precision plots (right) with 5-fold cross validation. (c) Top surface markers for identifying TM cells from non-TM cells in the blood based on COMET analysis (Delaney et al., 2019). The 20 top scoring genes by q-value are shown. Significance determined using an XL-minimal hypergeometric test with multiple hypothesis test corrections. Full list of markers available in Table S4. (d) Classification of biological functions for positive markers (q-value <= 0.01) identified for TM cells using COMET. (e) Frequency of CD8^+^ T cells in the blood of mice with MC38 tumors at day 21 post implantation (n=9 mice) that express the indicated proteins using flow cytometry. Cells are gated on singlets, live, CD45^+^, CD8α^+^. Data are representative of 2-4 independent experiments depending on the marker with n=5-9 mice. Bars show the mean, and error bars represent standard deviation. (f) Flow cytometry contour plots showing CD39, NKG2D, and CX3CR1 expression (Y axis) as indicated above each plot, and CD44 (X axis) on CD8^+^ T cells in the blood of mice in (e). (g) UMAP visualization of mice from a validation cohort (Mouse 4 and Mouse 5) combining gene expression, TCR, and surface protein expression using CITE-seq. Shown are CD8^+^ T cells colored by TM status in green (far left) or each of the indicated proteins (NKG2D, CD39, and CX3CR1). Red indicates positive expression of the protein. Grey indicates non-matching status in the far left UMAP and negative expression for the protein in the three UMAPs to the right. Significance for CD39: p=3.87×10^−54^ and p=7.53×10^−71^, NKG2D: p=3.19×10^−122^ and p=1.93×10^−175^, CX3CR1: p=9.22×10^−17^ and p=2.08×10^−30^ for Mouse 4 (M4) and Mouse 5 (M5) respectively, assessed using a Wilcoxon rank-sum test. (h) ROC curves showing the sensitivity and specificity of each protein (NKG2D, CD39, and CX3CR1) at identifying TM cells. (i) Scatter plot showing the sensitivity and specificity for NKG2D, CD39, and CX3CR1 protein expression in identifying TM cells, along with all two- and three-protein combinations of these three markers. Shown are both “and” combinations and “or” combinations, colored black if they are Pareto-optimal; that is, if there is no other gate with strictly better sensitivity *and* specificity. The symbol “&” indicates the “and” gate, and the “|” indicates the “or” gate. Points that are not Pareto-optimal are colored grey. Individual and select combinations are labeled. Full list of values is in Table S5.

To test whether single-gene markers could be used to identify the CD8^+^ TM component with flow cytometry, we applied COMET, a computational tool we previously developed to predict markers from single-cell RNA-seq data (Delaney et al., 2019). COMET identified 82 candidate positive markers for the CD8^+^ TM component (with q value <= 0.01) which we classified into four general biological categories: negative regulatory pathways (e.g. *Pdcd1, Lag3, Tigit*), positive regulatory pathways (including general activation, e.g. *Cd3e, Cd247*, costimulation, e.g. *Icos, Cd226, Slamf7*, and cytokine receptors/signaling, e.g. *Il18r1, Il18rap, Il2rb*), trafficking molecules (e.g. *Itga4, Itgb1, Cx3cr1, Ccr5*), and NK receptors (e.g. *Klrk1, Klrc1, Klrd1*) (Fig. 2C, 2D, Table S4). Additionally, COMET identified 21 candidate positive markers associated with non-TM cells (Fig. S2A), many of which were consistent with the naïve and/or central memory-like phenotype in non-TM cells (e.g. *Ccr7, Sell, Il7r*) (Fig. 1E, S1I).

Flow cytometry was used to validate that several markers detected at the transcript level were also detected at the protein level (Fig. 2E). Moreover, many of these markers were enriched on CD44^+^ cells (Fig. S2B). Some of these markers trended towards a higher frequency in the blood of mice bearing MC38 tumors than naïve mice, but many including PD-1 were not different (Fig. S2C). To test if surface proteins could be used to enrich for TM cells, we evaluated three of the COMET-predicted candidates (*Entpd1* encoding CD39, *Cx3cr1* encoding CX3CR1, and *Klrk1* encoding NKG2D) (Fig. 2C, 2D). Flow cytometric analysis showed that a small number of CD8^+^ T cells expressed each of these markers in the blood of mice with MC38 tumors (Fig. 2E, 2F), albeit less than observed in the tumor (Fig. S2D). We next conducted a single-cell validation experiment measuring simultaneously (1) gene expression, (2) TCR, and (3) protein expression for CD39, CX3CR1, and NKG2D using CITE-seq (Stoeckius et al., 2017) in two mice (Fig. 2G, S2E). We observed that each protein (NKG2D, CD39, and CX3CR1) could be successfully used to distinguish between CD8^+^ TM and non-TM cells and enrich for TM cells, and the relative ranking and overall performance of our three validated genes were consistent across mice (Fig. 2G, 2H).

We next asked whether NKG2D, CD39, and CX3CR1 were defining distinct subpopulations within the TM compartment, or improving on the technical accuracy of recovering a single, more homogenous population. Flow cytometry of NKG2D, CD39, and CX3CR1 in the bulk CD8^+^ T cell population showed various combinations of single, co-expression, or triple expression (Fig. S2F, S2G). To determine if co-expression of these three markers varied between the TM and non-TM components, we assessed their co-expression landscape in the CITE-seq data. Here, the frequency of cells expressing one, two, or three of the markers was comparable to the flow cytometry results when examining expression on all of the CD8^+^ T cells assayed (Fig. S2H). However, restricting to the TM cells using the TCR, the majority of the cells expressed two or three of the markers (Fig. S2I), suggesting that using these markers in combination could create more optimal balances between sensitivity and specificity. Indeed, using combinations of these markers improved on either or both the sensitivity and specificity as compared to using single markers (Fig. 2I, Table S5). For example, using the combination gate [NKG2D or CD39] improved the sensitivity to 0.94 from 0.85 or 0.62 (values are averages across the two mice) when using NKG2D or CD39 as singleton markers, respectively, while reducing the specificity. Similarly, the combination gate [NKG2D & CD39 & CX3CR1] enabled sorting with higher specificity than any of the single markers (0.98 on average), and hence achieved minimal contamination, while compromising on sensitivity (Fig. 2I). Consequently, while the TCR likely remains the most sensitive and specific metric for determining whether T cells have shared reactivity, cell surface markers can be used to distinguish TM cells from non-TM cells.

### Tumor-matching CD8^+^ T cells in blood are less dysfunctional than matching clones found in the tumor

We next examined the transcriptional heterogeneity of CD8^+^ T cells in the tumor whose TCRs were also detected in blood, referred to as “blood-matching” cells. In the tumor, a diverse range of differentiation states were observed (Fig. 1C, Table S1). Blood-matching cells were present in every transcriptional cluster in the tumor (Fig. 3A, 3B, S1D). The frequency of blood-matching cells varied across clusters and showed some mouse to mouse variability (Fig. 3A, 3B, S1D), but the majority of blood-matching cells were present in non-naïve/non-central memory-like clusters in the tumor (Fig. 3B), and were significantly more clonally expanded than non-matching cells (Fig. 3C, p=4.9×10^−26^). In Mouse 1, we observed a correlation between clone size in blood and clone size in tumor (Fig. 3D). While our clone sizes were too low in Mouse 2 and Mouse 3 to observe a significant correlation in expansion between blood and tumor, we did observe this correlation in the two mice in our validation cohort (Mouse 4 and Mouse 5) where the number of TM cells recovered was higher (Fig. S3A). To further characterize blood-matching T cells in the tumor, we examined signatures related to CD8^+^ T cell functions. We found that the blood-matching cells in the tumor expressed higher levels of a terminal exhaustion signature (Miller et al., 2019) compared to the non-matching cells (Fig S3B). Blood-matching cells also expressed higher levels of a TRM signature (Beura et al., 2018), associated with TRM cells which have been shown to play a role in protective immunity against tumors (Menares et al., 2019; Park et al., 2019) (Fig S3B). Blood-matching cells expressed lower levels of a naïve T cell signature (Kaech et al., 2002) compared to the non-matching cells, and no significant difference was observed in the extent of expression of a cell-cycle signature (Kowalczyk et al., 2015) (Fig S3B). Lastly, previous work has shown that pathogen-specific CD8^+^ T cells can infiltrate tumors in both mice and humans (Mognol et al., 2017; Rosato et al., 2019; Simoni et al., 2018), though the significance of these cells is unclear. This bystander signature (Mognol et al., 2017) was present in several of the transcriptional clusters within tumor (Fig. S3B), but was expressed at lower levels in blood-matching cells as compared to non-matching cells. These findings suggest that the TM component being detected in blood is corresponding to T cell populations in the tumor that are likely responding to tumor antigens, and are relevant for tumor killing.

**Figure 3:**
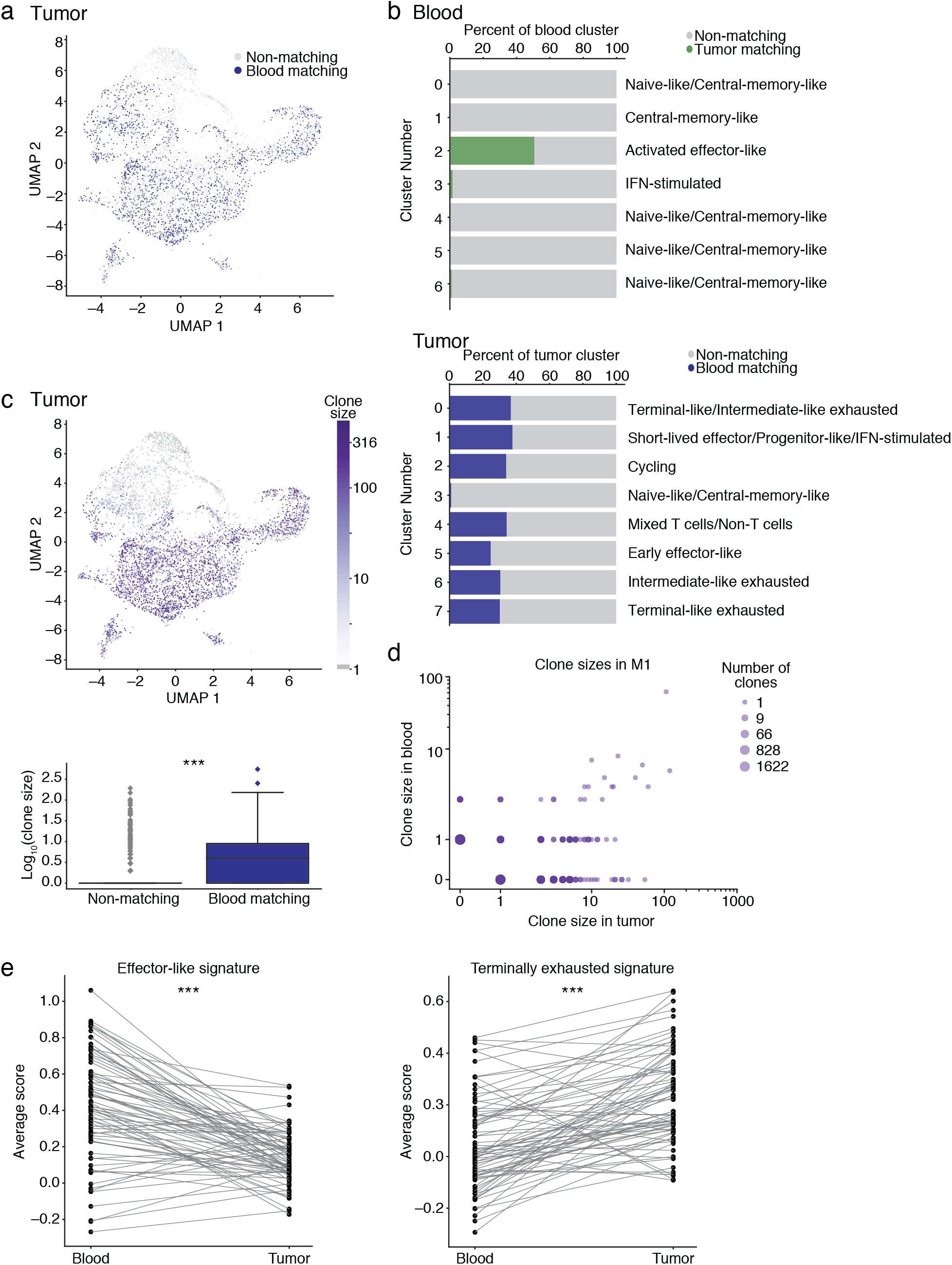
Tumor-matching CD8^+^ T cells in blood are less dysfunctional than the corresponding clones in tumor. (a) CD8^+^ T cells from the integrated MC38 tumor samples that share a TCR identified in the paired blood sample are indicated in navy blue (referred to as blood-matching cells). Non-matching cells are indicated in grey. (b) The distribution across identified transcriptional clusters of TCR matched cells between tumor and blood. Shown is the percentage of each cluster that is matching vs. non-matching. Shown are clusters with more than 50 cells. (c) UMAP visualization showing clonal expansion across CD8^+^ T cells from tumor as measured by clone size (Top). Box plot quantifying the amount of clonal expansion in the tumor (Bottom). Significance determined using the Wilcoxon rank-sum test. p=4.9×10^−26^. (d) Expansion rates of clones in blood and MC38 tumor (log-scale, for M1). (e) Cells within the same clone have a higher “terminal exhaustion” gene signature (p=3.1×10^−9^) and a lower “effector-like” gene signature (p=1.1×10^−9^) (signatures from (Miller et al., 2019)) in tumor compared to blood. Significance determined using a Wilcoxon signed-rank test. Each dot shows the average gene signature of the cells in a given clone, and lines connect the same clone between blood and tumor samples. Shown are clones detected in M1. M2 and M3 shown in Fig. S3D.

Next, we compared the transcriptional profiles of TM cells in blood to their matching clones in the tumor (the blood-matching compartment). The transcriptional profiles of blood-matching cells within tumor were more diverse than the transcriptional profiles of the TM cells in blood, in that the TM cells in blood were primarily found in one transcriptional cluster, whereas their counterparts in tumor where found in the large majority of clusters (Fig. 3B). This observation suggests that upon entering the tumor, CD8^+^ T cells can diversify and take on a number of different phenotypes. We found that the blood-matching component within tumor expressed higher levels of a terminally exhausted signature (Miller et al., 2019) as compared to the TM cells in blood, and that the TM cells in the blood expressed higher levels of an effector-like CD8^+^ T cell signature (Miller et al., 2019) as compared to blood-matching cells in tumor (Fig. S3C). These data are consistent with the cells in the tumor being more dysfunctional than the cells in the blood. Next, we performed differential gene expression analysis on clonally matched populations between the blood and tumor compartments, considering each matched clone in tumor or blood as a single observation (see Methods). Key genes up regulated in TM clones in the blood compared to blood-matching clones in tumor included *Ccl5, Cx3cr1, Itga4, Runx1,* and *Klrg1* (Table S6), consistent with an effector-like state of these cells. Key genes up regulated in the blood-matching clones in tumor included *Pdcd1, Lag3, Ctla4, Havcr2,* and *Tigit* (Table S6), again consistent with the blood-matching clones in the tumor being more dysfunctional than matching cells in blood. The blood-matching clones in tumors also showed elevated levels of many of the granzymes (*Gzmb, Gzmc, Gzmd, Gzme, Gzmf,* and *Gzmg*) compared with the TM clones in blood (Table S6), consistent with previous work showing some overlap between effector-associated genes and exhausted T cell populations, particularly terminally exhausted T cells (Beltra et al., 2020; Singer et al., 2016).

Lastly, we examined the transcriptional state of CD8^+^ T cells in the two tissue compartments on a clone by clone basis. This enabled us to distinguish whether the transcriptional state of a clone is determined by the TCR alone, or whether the transcriptional state of cells is different across blood and tumor in cells with the same TCR. We found that the cells within a clone in blood were significantly more enriched for an effector-like signature than the cells within the matching clone in tumor (M1 p=1.4×10^−9^, M2 p=0.0072, M3 p=0.036), and that cells within a clone in the tumor were significantly more enriched for the terminal exhaustion signature than the cells within the matching clone in blood (M1 p=2.6×10^−9^, M2 p=0.00053, M3 p=0.0011) (Fig. 3E, S3D) (signatures from (Miller et al., 2019)). These data suggest that for individual clones, the TM cells in blood are more effector-like than cells in the tumor, and that after migration into the tumor these cells acquire a dysfunctional state.

### Activated tumor-matching CD8^+^ T cells can be detected in the blood of melanoma patients

Given our ability to identify TM CD8^+^ T cells in the blood of mice with MC38 tumors, we next asked if this approach could be used in cancer patients. We used coupled single-cell RNA-seq and TCR sequencing from paired blood and tumor samples obtained from four checkpoint treatment-naïve advanced melanoma patients (Fig. S4A-S4H, Table S7). Here, “tumor” refers to tissue resections obtained from either the primary tumor site or metastases (lymph node or subcutaneous) as indicated for each patient (Fig. S4B, Table S7). Characterization of the CD8^+^ T cells across patients revealed transcriptional signatures in blood consistent with naïve-like, central memory-like, effector-like, and effector memory-like cells, and signatures in tumor consistent with diverse exhausted subpopulations, effector-like, resident memory-like, naïve-like and/or central-memory like, and cycling populations, as previously reported in the literature (Guo et al., 2018; Sade-Feldman et al., 2018; Siddiqui et al., 2019; Tirosh et al., 2016; van der Leun et al., 2020; Yost et al., 2019) (Fig. 4A, 4B, Table S8).

**Figure 4:**
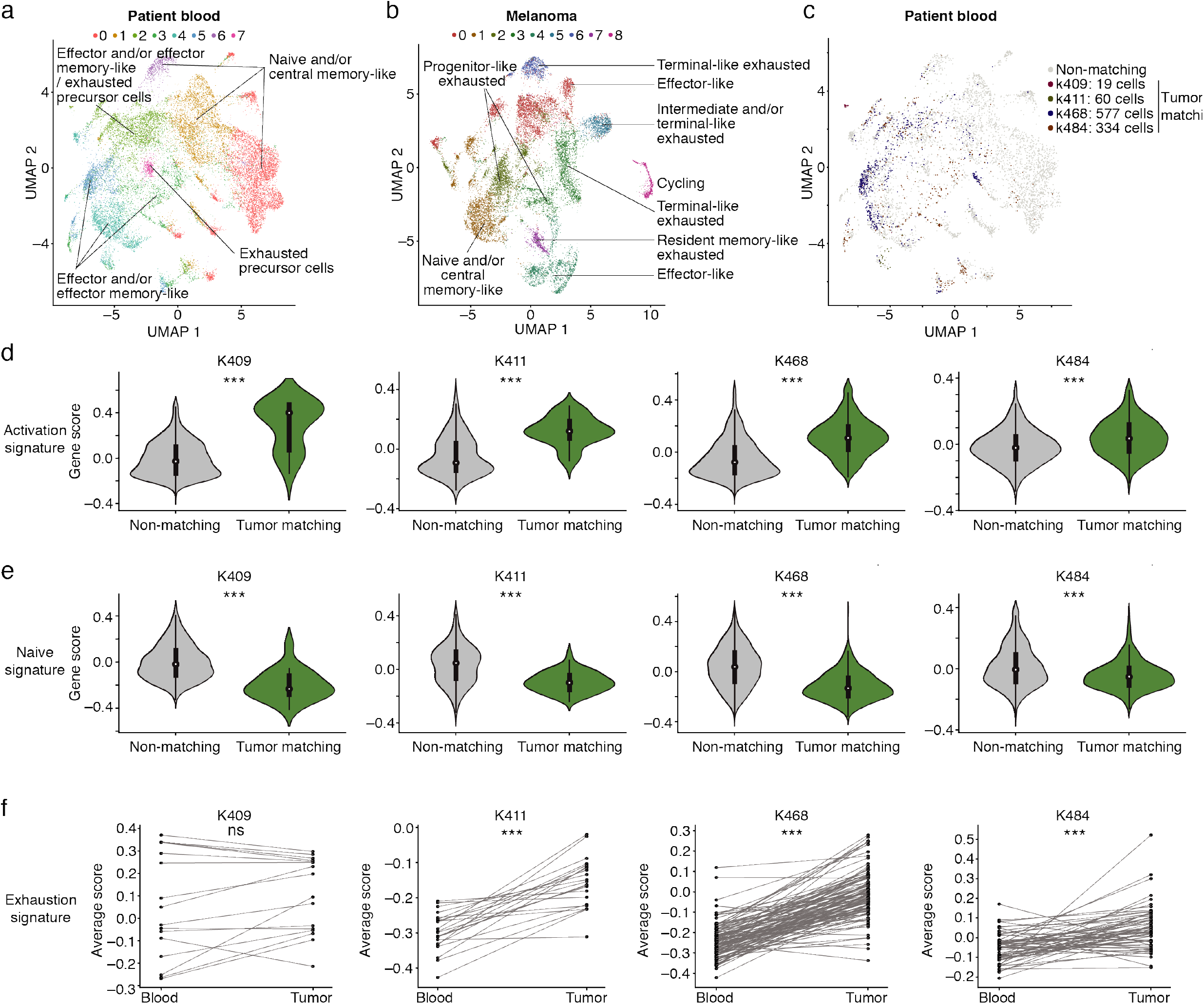
Tumor-matching CD8^+^ T cell clones can be detected in the blood of metastatic melanoma patients and show less signs of dysfunction than matching clones in tumor. (a-b) Clustering and UMAP visualization of paired blood (n= 21,833 cells) and tumor (n= 16,878 cells) samples from immunotherapy treatment naïve patients. Data are integrated from the initial blood sample from four patients (patient clinical parameters shown in Fig. S4A and Table S7). Colors indicate transcriptional clusters. Labels marking each cluster indicate the phenotypic description of the cluster based on gene expression data (see Methods). Up regulated genes in each cluster are in Table S8. Cells displayed filtered to show CD8^+^ T cells. (c) CD8^+^ T cells in blood colored in the integrated human UMAP that have a TCR matching to CD8^+^ T cells found in tumor (TM cells), colored by each patient. Grey indicates CD8^+^ T cells that do not have a TCR matching to T cells in tumor (referred to as non-TM). (d-e) Violin plots showing enrichment of activation (d) or naïve (e) CD8^+^ T cell signatures in TM and non-TM cells. Signatures derived from (Akondy et al., 2017). Significance determined using a Wilcoxon rank-sum test. For the activation signature in (d), p values are K409 p=7.1×10^−8^, K411 p=3.3×10^−15^, K468 p=3.2×10^−101^, K484 p=6×10^−15^. For the naïve signature in (e), p values are K409 p=1.1×10^−7^, K411 p=8.9×10^−11^, K468 p=1×10^−91^, K484 p=2.9×10^−12^. (f) Mean value of an “exhaustion” signature from (Sade-Feldman et al., 2018) on a clone by clone basis in blood and in tumor. Significance determined using a Wilcoxon signed-rank test, p values are K409 p=0.2, K411 p=4×10^−5^, K468 p=6.5×10^−19^, K484 p=2.1×10^−7^. Each dot shows a clone, and lines connect the same clone between paired blood and tumor samples. For patient samples, “tumor” in the Figure refers to both resections from the primary tumor and metastases as indicated in Fig. S4B.

We next determined whether we could identify TM cells in patients by using the TCR as a molecular barcode. We detected TM cells in the blood samples of each of the four patients, with the number of cells recovered varying by patient (Fig. 4C, S4B). Despite the heterogeneity observed across patients (Fig. S4E, S4F), the TM cells were generally observed in regions of the UMAP associated with activated T cell phenotypes (Fig. 4A, 4C). Indeed, the majority of TM cells in each patient were present in “non-naïve” clusters (e.g. not clusters 0, 1, or 6, which were associated with naïve-like and/or central-memory like cells). The percentages of TM cells in these “non-naïve” clusters were 94.7% (K409), 88.3% (K411), 83.5% (K468) and 87.7% (K484). Many of the TM cells belonged to clusters associated with an effector-like and/or effector memory-like phenotype (clusters 2, 3, 4, and 5). Consistent with this, TM cells in the blood of all four patients expressed significantly higher levels of an activation signature compared to non-TM cells (Fig. 4D), and non-TM cells expressed significantly higher levels of the naïve signature compared to TM cells (Fig. 4E) (signatures derived from (Akondy et al., 2017)). To interrogate how the level of exhaustion compared between clones in blood and clones in tumor, we evaluated the enrichment of an exhaustion signature (Sade-Feldman et al., 2018) on a clone by clone basis between these two tissues. In patient K409, there was no significant difference in the exhaustion score between clones in blood and tumor (Fig. 4F, K409 p=0.2), suggesting that the clones in the blood of this patient may be exhausted. However, in the other three patients analyzed, we did observe an increase in this signature of exhaustion (Sade-Feldman et al., 2018) on matching clones in tumor relative to blood (Fig. 4F, K411 p=4×10^−5^, K468 p=7.1×10^−19^, K484 p=1.9×10^−7^). These data are consistent with our results in mice, supporting the idea that TM cells in the blood may be less dysfunctional than their corresponding counterparts in tumor despite shared TCR sequences.

### Tumor-matching CD8^+^ T cells can be tracked longitudinally in patient blood and show a temporal increase in exhaustion despite anti-PD-1 treatment

In addition to the four initial treatment naïve samples, longitudinal blood samples were collected from two patients that went on to fail to respond to checkpoint blockade, K411 and K468 (Fig. S4A). Both patients had tumor at the time of the longitudinal sample collection (Fig. S4A). We first determined whether TM cells could be identified in the follow up blood sample based on TCR sequence. Here, we detected overlapping clones (TM cells) between the two blood samples and the tumor sample in each patient, despite one of the samples being collected almost a year and half after the initial sample (Fig. 5A, S4A). The TM cells detected in the longitudinal samples showed signs of activation compared to non-TM cells (Fig. S4I, S4J), similar to the trend in the initial sample (Fig. 4D, 4E). Interestingly, when comparing the signature of exhaustion (Sade-Feldman et al., 2018) between the three samples (tumor, treatment naïve blood, and longitudinal blood), we observed that the exhaustion signature in the longitudinal samples was higher than the initial blood sample, but lower than the tumor (Fig. 5B). These data suggest that the TM population in the blood can become more exhausted over time despite anti-PD-1 treatment, but that ultimately the highest levels of exhausted T cells remain in the tumor microenvironment (Fig. 5B).

**Figure 5:**
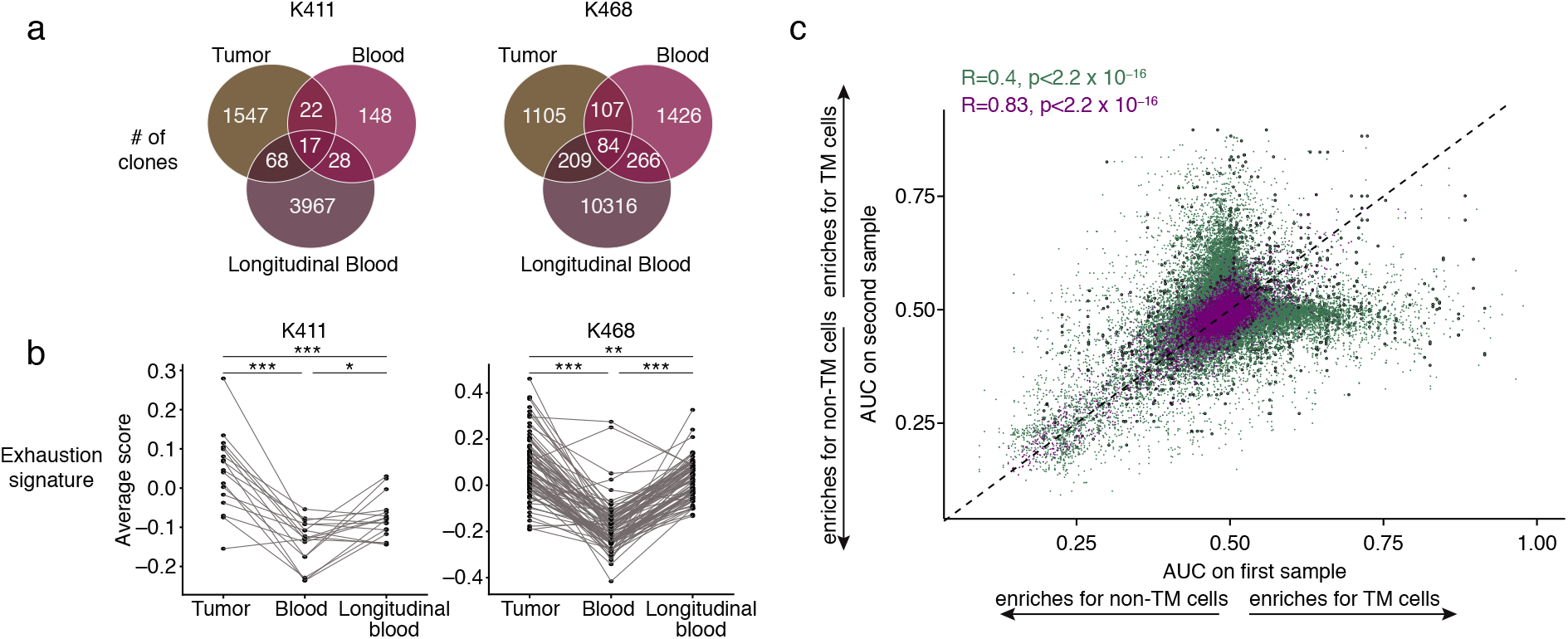
Matching clones can be detected in longitudinal blood samples from melanoma patients and show increased signs of exhaustion compared to clones from pre-treatment blood. (a) Number of clones detected and overlapping between samples in the initial blood and tumor sample of K411 and K468, as well as the longitudinal blood sample collected (patient clinical parameters shown in Fig. S4A and Table S7). (b) Mean value of an “exhaustion” gene signature from (Sade-Feldman et al., 2018) on a clone by clone basis in tumor, initial paired blood, and longitudinal blood. Each dot shows a clone, and lines connect the same clone between the original paired blood and tumor samples, as well as the matched longitudinal blood sample. Analyzed are only clones that were detectable in both blood samples and the tumor sample. Significance determined using a Wilcoxon signed-rank test. For patient K411, p=0.022 between blood and longitudinal blood, p=3.5×10^−4^ between blood and tumor, p=8.5×10^−4^ between longitudinal blood and tumor. For patient K468, p=1.1×10^−14^ between blood and longitudinal blood, p=5.8×10^−15^ between blood and tumor, p=0.0017 between longitudinal blood and tumor. (c) Scatter plot showing the degree to which each gene’s AUC for selecting TM cells from blood at the transcriptional is concordant across patient samples. Purple denotes a comparison between longitudinal samples from the same patient; Green denotes a comparison between different patients. Points outlined in black are surface-expressed genes. Significance determined using the Spearman correlation test. For patient samples, “tumor” in the Figure refers to both resections from the primary tumor and metastases as indicated in Fig. S4B.

Having observed significant longitudinal shifts in the transcriptional state of the TM component, we next quantified the extent to which the transcripts enriched in the TM component relative to the non-TM component correlated across patient samples. We observed that the extent of similarity across samples is greater for within-patient comparisons (longitudinal samples generated from the same patient) than between-patient comparisons (Fig. 5C, Table S9). Notably, despite the acquired differences in the T cell exhaustion signature of clones following therapeutic intervention (Fig 5B), the general transcriptional landscape of the TM component relative to the non-TM component remained highly consistent within the two patients assessed in this study (R=0.83, p-value < 2.2×10^−16^).

Analysis of between-patient variability revealed a significant correlation (Fig. 5C, R=0.4, p-value < 2.2×10^−16^) in the extent to which individual gene transcripts are specific to the TM component or the non-TM component. This consistency across patients suggested that there may exist useful transcripts for isolating the TM component from blood. Since these transcripts would have a number of practical uses if expressed on the cell surface (e.g. sorting for sequencing, functional assays, or adoptive cell transfer therapy), we restricted our correlation analysis to a gene set of cell surface markers (Chihara et al., 2018) and the significant correlations in the TM component remained (R=0.31, p value < 2.2×10^−16^). The highly significant correlation of surface-expressed, TM-enriched transcripts across patients and therapeutic states suggests that general biomarkers could be defined for the TM population that are robust to varying tumor burdens and therapeutic conditions.

### Transcriptional analysis suggests that cell surface marker combinations can be used to detect the TM component from patient blood

Having identified a TM component in patient blood with distinct transcriptional properties compared to non-TM cells, we asked whether cell surface markers could be used to sort the TM component from the non-TM component. We first examined use of inhibitory receptors to identify TM cells, since markers such as PD-1 have been used to study the putative immune response to tumor in peripheral blood (Gros et al., 2016; Gros et al., 2014; Huang et al., 2019; Huang et al., 2017; Kamphorst et al., 2017). We observed that with the exception of patient K409, *PDCD1* RNA was detected only on a minority of the TM cells (Fig. 6A). Moreover, at the transcript level, *PDCD1* and a number of other inhibitory receptors (*HAVCR2, LAG3, CD160, TIGIT,* and *BTLA*) had very poor performance (nearly indistinguishable from random) as predictive markers across the range of potential threshold values in both the initial blood sample (Fig. 6B, Table S9) and longitudinal samples (Fig. 6C, Table S9). Our finding that the AUC for the inhibitory receptors is hardly above 0.5 for most patients suggests that this class of markers cannot be relied upon to enrich for the TM component of patient blood.

**Figure 6:**
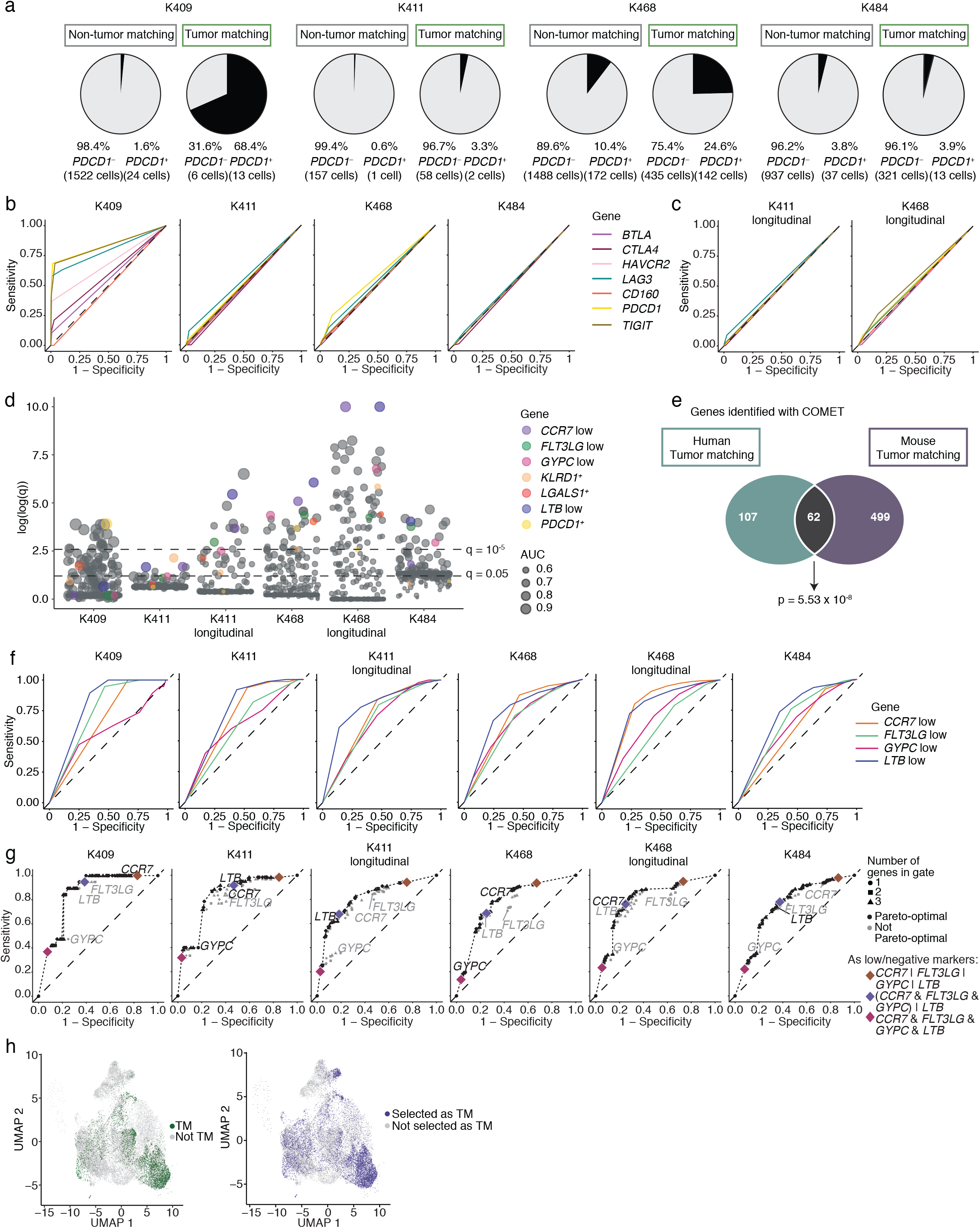
Identification of combinations of markers for tracking tumor-matching clones across patients. (a) Frequency of *PDCD1*^+^ cells (using transcript) in the original blood sample separated by TM and non-TM cells. (b-c) ROC curves showing the sensitivity and specificity of *PDCD1, BTLA, CTLA4, HAVCR2, LAG3, CD160,* or *TIGIT* to distinguish TM cells from non-TM cells in (b) the original blood samples and (c) in the longitudinal blood samples from K468 and K411 at the transcript level. Legend indicating colors shared between (b) and (c). (d) Plot showing the significance values from the COMET analysis across patient samples. Significance determined using an XL-minimal hypergeometric test with multiple hypothesis test corrections. Circles are sized by the AUC for that gene in sorting TM cells from non-TM cells. The y axis corresponds to the log2(x+1) transformation of the −log10 of the COMET q values, capped at 10. Consensus markers are highlighted with color, along with *PDCD1* for reference. All other surface-expressed markers shown in grey. Full list of markers in Table S10. (e) Overlap between the single-gene markers detected by COMET to distinguish TM cells from non-TM cells in mouse (MC38 tumors) and human (melanoma patients) samples. Single-gene markers were included if detected as significant by COMET (q-value <0.05) in a minimum of two samples (see Table S11 for overlap tests across additional parameters). Significance determined using a hypergeometric test, p=5.53×10^−8^. (f) ROC curves for the consensus markers identified in (d) for each patient. (g) The sensitivity and specificity of all possible logic gates derived from combinations of genes *CCR7*^low^, *LTB* ^low^, *GYPC* ^low^, and *FLT3LG* ^low^ at their COMET-determined cutoff values (available in Table S10). Points are shaped by the number of markers used in the logical gate, and colored black if they are Pareto-optimal (if there is no gate with strictly better sensitivity *and* specificity). A dotted line through the Pareto-optimal gates represents the ROC of this combinatorial marker collection. For (f) and (g), the dashed line (on the diagonal) represents the sensitivity and specificity values of random chance. (h) UMAP visualizations of CD8^+^ cells integrated from all patient blood samples (including longitudinal samples). Left: true tumor-matching cells as defined by matching TCR sequence are colored green. Right: putative tumor-matching cells as determined by the best-performing gate, [*(CCR7* ^low^ & *FLT3LG* ^low^ & *GYPC*^low^) | *LTB*^low^], are colored blue. For this combination, Sensitivity = 0.751, Specificity = 0.745. The symbol “&” indicates the “and” gate, and the “|” indicates the “or” gate.

To determine if better markers for isolating these cells could be identified in humans as we had done in mice, we again used COMET (Delaney et al., 2019) to evaluate surface-expressed genes (Chihara et al., 2018). For each of the six patient blood samples, COMET identified transcripts that significantly enriched for the TM component (Fig. 6D, Table S10). Interestingly, we observed a significant overlap between candidate markers for the TM compartment identified from the patient samples and the markers identified in mice (Fig. 6E and Table S11), suggesting that some markers of TM cells are conserved across species and cancer types. We identified 15 near-consensus surface markers that had q < 0.05 in at least four of the six patient samples (Table S12) as candidates for sorting. Of the 15 near-consensus genes, many had lower expression in TM cells than in non-TM cells (henceforth referred to as “negation markers”). The top four ranking genes based on AUC for distinguishing TM cells from non-TM (see Methods), *LTB, CCR7*, *GYPC,* and *FLT3LG* were negation markers (referred to as *LTB*^low^*, CCR7*^low^*, GYPC*^low^*, and FLT3LG*^low^). Low expression of these markers is consistent with the non-naïve, largely effector or effector-memory like transcriptional state of many of the TM cells (Fig. 4A, 4C). These four markers showed consensus across patients with respect to AUC performance, despite differing tumor burdens and therapeutic states (Fig. 6D, 6F, Table S13). Indeed, these four markers have significantly high average AUC performance across all patient samples (Fig. S4K). These four consensus markers featured differing strengths in sensitivity in specificity (Table S13, see Methods); for example, *CCR7*^low^ achieved 0.827 sensitivity and 0.621 specificity (empirical p < 0.0001), while *GYPC*^low^ achieved 0.340 sensitivity and 0.819 specificity (empirical p < 0.0001). *FLT3LG*^low^ and *LTB*^low^ each exhibited a more even balance between sensitivity and specificity, 0.780 and 0.447 for *FLT3LG*^low^, respectively, and 0.718 and 0.768 (for *LTB*^low^, respectively (empirical p < 0.0001 for each, Fig. 6G, Fig. S4L, Table S13, see Methods). When evaluated through a range of expression cutoff values, these markers demonstrated remarkable AUC performance that was robust to sampling variation (*CCR7*^low^: 0.742; *FLT3LG*^low^: 0.619; *GYPC*^low^: 0.647; *LTB*^low^: 0.772; empirical p < 0.0001 for each; Table S13, see Methods). Though all four of these consensus markers are negation markers (meaning their expression is lower in TM cells), we did observe some positive markers for TM cells lower on the ranked list, including *KLRD1* and *LGALS1*, which came up in a companion study co-submitted with this work (Lucca et al., submitted) (Fig. 6D and Table S12).

To increase performance of a sorting strategy for TM cells from blood, we next explored the use of combinations of cell surface markers. We evaluated the performance of the four top-ranking markers (*LTB*^low^*, CCR7*^low^, *GYPC*^low^, and *FLT3LG*^low^) as well as all possible two, three, and four marker logical gates derived from these markers (Fig. 6G-H, Table S14, see Methods). We observed that in all samples, all four markers enrich for the TM component (as negation markers), and that marker combinations of two or more genes significantly improved performance on the sensitivity (capture rate) and/or specificity (contamination rate) axes. The best-performing gate with even balance between capture rate and contamination rate, [*(CCR7*^low^ *and FLT3LG*^low^ *and GYPC*^low^*) or LTB*^low^] (meaning that a TM cell has low expression of LTB, or has low expression of any of the other three markers), isolated the TM component of blood pooled across all patient samples with 0.751 sensitivity and 0.745 specificity (empirical p < 0.0001 for each) and is consistent in its performance in individual patients (Fig. 6G-H, Table S13, see Methods). Importantly, marker combinations of these four genes enabled achieving a broad range of sensitivity and specificity, and many of the marker combinations showed consistency in their sensitivity and specificity rates across patients (Fig 6G, Table S13 and S14, see Methods). Collectively, these data highlight the utility in using combinations of markers, and demonstrate how the sensitivity and specificity can be tuned depending on the goal of the experiment.

Lastly, we determined how robust these marker combinations would be if we used alternative methods of defining matching clones between blood and tumor. Using an exact sequence match for both the alpha and beta chain (see Methods) is a highly stringent definition, and it is possible that some TM cells may have been missed. To address this issue, we utilized two additional TCR clustering tools, GLIPH2 (Huang et al., 2020) and iSMART (Zhang et al., 2020) to identify tumor-matching cells in blood. Across all patients, we observed an average increase of 8.2% in the number of TM cells (5.26% - 12.9%) (Fig. S5A). However, we still observed a significant enrichment in an activation signature in the TM cells relative to the non-TM cells in each patient (Fig. S5B), and a significant enrichment in the naïve signature in the non-TM cells (Fig. S5C) (signatures derived from (Akondy et al., 2017)). Additionally, the sensitivity of *PDCD1* and the other inhibitory receptors remained insufficient overall (Fig. S5D, S5E, Table S15). In contrast, the AUC performance of *CCR7*^low^*, FLT3LG*^low^*, GYPC*^low^, and *LTB*^low^ remained high across all of the patient samples (Fig. S5F, Table S15). While future studies coupling larger cohorts with CITE seq will be important to determine how generalizable these findings are across patients and to validate key markers, the concept that surface marker panels for the TM component could be built from these types of analyses to monitor patients’ response to immunotherapy in real time has tremendous clinical potential.

## Discussion

With the rapidly increasing use of immunotherapy to treat cancer, there is tremendous interest in monitoring the anti-tumor immune response. However, the inability to routinely identify the tumor-relevant CD8^+^ T cell component in blood limits such monitoring. In our study, we used the TCR as a molecular barcode as a method to identify TM CD8^+^ T cells in the blood of mice with MC38 tumors and melanoma patients. TM cells showed increased signs of activation compared to non-TM cells in the blood, but were less dysfunctional than their counterparts in tumor. We applied an algorithmic approach to identify candidate marker panels from the transcriptional profile of TM cells. Using CITE seq, we validated that three of these candidate markers at the protein level (NKG2D, CD39, and CX3CR1) successfully identified the TM component from the non-TM component in mice, and that marker combinations provided a broad range of sensitivity and specificity values when isolating the TM component. Collectively, our data highlight the utility of using the TCR to identify TM cells in blood, enabling the generation of marker panels to aid in the isolation of tumor-relevant populations for downstream applications.

The blood is a conduit of immune cell trafficking, acting as a highway between secondary lymphoid organs and non-lymphoid tissues including tumors (Masopust and Schenkel, 2013; Mueller et al., 2013). As such, the blood has long served as a window into host immune responses. However, comprehensive profiling of tumor antigen-specific populations in the blood has been challenging. Use of the TCR as a molecular barcode to track TM cells provides an effective, less biased approach for identifying “tumor relevant” populations than alternative methods like PD-1 expression. Additionally, use of TCR sequencing captures a larger breadth of the anti-tumor response than individual peptide/MHC tetramers. Despite these notable advantages, there are technical and biological considerations associated with this method. First, paired blood and tumor samples are required from the same patient to identify TM cells. Second, it is possible that the entire repertoire of TM cells found within the blood may not be detected due to sampling depth issues in the tumor. Here, despite variability in the depth of tumor TCR coverage from patient to patient, these true TM cells showed consistent transcriptional states and markers that could be used for their isolation, highlighting the utility of the TCR to identify these cells. Additionally, when we used TCR clustering as a less stringent method to identify shared clones than exact sequence matching (GLIPH2 and iSMART), the sensitivity of *PDCD1* and the other inhibitory receptors remained poor, while the AUC performance of *CCR7*^low^*, FLT3LG*^low^*, GYPC*^low^, and *LTB*^low^ remained high. Third, use of negation markers can be challenging in single cell data, since these data sets contain a large number of zero values and it is has been debated whether counts of zero are due to true biology or technical artifacts. It is generally accepted that genes receiving zero counts are either not expressed or expressed to a low extent within a cell (Choi et al., 2020), and recent work has concluded that the zero measurements in count data reflect true biology (Choi et al., 2020; Hafemeister and Satija, 2019; Svensson, 2020; Townes et al., 2019). We therefore conclude, from the highly significant proportion of zeros and low count measurements in TM cells compared to non-TM cells as detected by COMET, that TM cells are lower for *GYPC, CCR7, LTB, and FLT3LG* than non-TM cells. Fourth, bystander T cells specific for pathogens have been identified in both mouse and human tumors using either peptide/MHC tetramers or TCR transgenic T cells (Mognol et al., 2017; Simoni et al., 2018). Here we observed evidence of a transcriptional signature of bystander cells in some of the tumor clusters, but the blood-matching component showed a lower enrichment than the non-matching component for this bystander signature in our samples. Hence, bystander cells were not a significant component of the blood-matching cells within tumor. Further work is needed to determine how many of the cells in the tumor with the bystander transcriptional signature indeed have TCRs that are specific to only to pathogens and not tumor, especially since it is possible that there could be some shared epitopes between pathogens and tumors. Follow up studies aimed at developing marker panels that can distinguish between these tumor-specific and bystander cells will be useful for immune monitoring purposes in cancer patients.

Here we found that TM cells differed in their transcriptional state compared to non-TM cells in the blood, showing increased signs of activation and effector function. Some of the markers of TM cells may be specific to this particular type of tumor or its location in the skin, and may differ with tumor type or location. It also should be noted, however, that while some of the trafficking molecules detected are associated with skin homing (including *Selplg*), a number are associated with a general program of trafficking to inflamed tissue sites and are not skin specific, including *Ccr2, Ccr5, Cx3cr1, Itga4, Itgb1,* and *Itgb2* (Liu et al., 2006; Masopust et al., 2010). Moreover, our observation that there is significant overlap between the genes identified in the mouse MC38 model and melanoma patients (Fig. 6E and Table S11) shows that there are similarities that span tumor type and species.

Interestingly, while a number of inhibitory receptors were detected at the transcript level, many were not detected at the protein level using flow cytometry, and as a class these transcripts performed poorly at distinguishing TM cells from non-TM cells in the blood. These results argue that there are better cell surface markers than the inhibitory receptors for identifying putative tumor-relevant populations in the blood, and highlight the utility of using the TCR as a way to enrich for the TM population for more comprehensive characterization. Three markers validated here for the identification of TM cells were NKG2D, CD39, and CX3CR1. When comparing effector, memory, and exhausted populations, *Klrk1* shows the highest expression in memory CD8^+^ T cells (assessed from GSE41867, (Doering et al., 2012)), and previous work has shown that NKG2D is important for optimal memory formation (Andre et al., 2012; Ferrari de Andrade et al., 2018; Prajapati et al., 2018; Wensveen et al., 2013; Zloza et al., 2012). CD39 has been associated with exhaustion in both mice and humans (Gupta et al., 2015). Levels of CX3CR1 have been shown to correlate with effector CD8^+^ T cell differentiation, with the highest levels of CX3CR1 being associated with the most effector-like cells (Gerlach et al., 2016). How NKG2D, CD39, and CX3CR1 as well as the other markers identified impact the function of TM cells remains to be determined.

In mice, we found that the population of TM cells in the blood was fairly homogenous, being largely enriched in one transcriptional cluster. However, the blood matching clones found within tumors showed significant transcriptional diversity, being detected within all of the different transcriptional clusters. These data suggest that TM CD8^+^ T cells have a high degree of plasticity in terms of differentiation upon entering the tissue, and the tumor microenvironment influences the development of a number of functional states. Our clone by clone analysis showed that on a clonal basis, TM cells in the blood were less exhausted than their blood-matching counterparts in tumor. Interestingly, for our two longitudinal patient samples, we observed that the clones detected in the second blood sample were more dysfunctional than the first, consistent with the notion that exhaustion continues to develop over time (Wherry and Kurachi, 2015). Ultimately, however, clones found in both the initial and later blood sample showed less of an enrichment in the exhaustion signature than the corresponding clones in the tumor, suggesting that relative to the tumor, the blood may still represent a reservoir of less dysfunctional cells.

In summary, by pairing single cell RNA sequencing with TCR analysis, we identified populations of CD8^+^ T cells in blood that had matching TCRs with CD8^+^ T cells in both mouse and human tumors. Using the TCR as a molecular barcode, we characterized the TM compartment in blood, and found that these cells were less dysfunctional than matching cells in the tumor. In addition, we provide robust evidence for an exciting and tractable innovation: the use of combinatorial marker panels to isolate the TM population in the blood. The combinatorial marker panels we identify are consistent over time, across patients, and robust to sampling variation. Follow up studies interrogating how immunotherapy strategies such as PD-1 blockade impact the TM population in the blood relative to the tumor as well as the overall clonality of the matching populations will be highly relevant to determining predictors of response versus resistance. Our algorithmic approach to generate marker panels to identify TM CD8^+^ T cells coupled with future longitudinal studies could assist with creation of diagnostics that aim to predict response to immunotherapy, potentially allowing monitoring of the anti-tumor immune response in patients in real time without the need for single-cell sequencing.

## Author Contributions

K.E.P., J.M. Luber, M.D.R, A.I.D, A.H.S. and M.S. conceived the study. K.E.P., J.M. Long, M.E.F., S.G., and K.P.B. performed mouse experiments. K.M.M., M.M.L., K.K.T., M.C., M.D.R., and A.I.D. participated in patient care, sample acquisition/coordination, and/or performing experiments with patient samples. K.A.L., O.S., J.M. Luber, L.H., C.D., K.N. and MS performed computational analyses. K.E.P., K.A.L., O.S., K.M.M., J.M. Luber, M.M.L., L.H., C.D., J.R.K., J.M.S., M.D.R, A.I.D, A.H.S. and M.S. contributed to data interpretation. K.E.P., K.A.L., O.S., K.M.M., J.M. Luber, M.M.L., M.D.R, A.I.D, A.H.S. and M.S. contributed to writing the manuscript. All authors reviewed the manuscript. M.D.R, A.I.D, A.H.S. and M.S. supervised the study.

## Acknowledgements

We would like to thank Christina Usher for preparation of the figures. We acknowledge the Parnassus Flow Cytometry Core at the University of California San Francisco supported in part by Grant NIH P30 DK063720 and by the NIH S10 Instrumentation Grant S10 1S10OD021822-01 for human samples, and the Immunology Research Flow Cytometry Core Facility at Harvard Medical School for performing FACS-based sorting for mouse samples. We would like to thank Dr. Seth Maleri for assistance with mouse single-cell library preparation. This manuscript was supported in part by the Parker Institute for Cancer Immunotherapy as well as grants from the National Institute of Health R01CA229851, U54CA224088, and PO1 AI56299 (to A.H.S.) and PO1 AI39671 (to A.H.S. and M.S.). Additionally, individual authors were supported by the following sources: T32GM007753 from the National Institute of General Medical Sciences (to K.A.L.) (the content is solely the responsibility of the authors and does not necessarily represent the official views of the National Institute of General Medical Sciences or the National Institutes of Health), NIH T32 Training Grant in Gastrointestinal Surgery (5T32DK007573) to (K.M.M.), National Science Foundation Graduate Research Fellowship to (J.M. Luber), (M.M.L.) was supported in part by an Abbvie Sponsored Research Award, the National Institute of General Medical Sciences Award T32GM007753 (to J.R.K.), and T32 CA009172 and as a Janock Fellow in Gastrointestinal Malignancies at the Dana Farber Cancer Center (to K.P.B).

The authors declare the enclosed potential conflicts of interest. M.D.R. is a founder of TRex Bio and Sitryx Bio and receives research funding from Abbvie, LEO Pharma, and TRex bio. A.I.D. has funds from Merck, Oncosec, BMS, Roche, Genentech, Pfizer, Incyte, Novartis and Checkmate, is on advisory boards for Xencor, Pfizer, Array, has stock in TRex, SQZ bio., and patents with Oncosec on Gene Therapy of melanoma. A.H.S. has patents on the PD-1 pathway licensed by Roche/Genentech and Novartis, consults for Novartis, is on the scientific advisory boards for Surface Oncology, Sqz Biotech, Elstar Therapeutics, Elpiscience, Selecta and Monopteros, and has research funding from Merck, Novartis, Roche, and Quark Ventures. K.K.T discloses institutional research funding from Array/Pfizer, BMS, Oncosec, Regeneron, and Replimune. From August 4^th^ 2020, MS is an employee of Guardant Health. The authors have no additional financial conflicts of interest to disclose.

## Materials and Methods

### EXPERIMENTAL MODEL AND SUBJECT DETAILS

#### Mice and Cell Lines

Wild type (WT) female C57BL/6 mice were purchased from the Jackson Laboratory (stock number 000664). Tumor cells were implanted into mice at 8-10 weeks of age. Mice were maintained at Harvard Medical School in specific pathogen-free facilities under standard housing, husbandry, and diet conditions in accordance with Institutional Animal Care and Use Committee (IACUC) and NIH guidelines. All experimental procedures performed were approved by the IACUC at Harvard Medical School.

For tumor studies, MC38 colon adenocarcinoma cells (a gift from Dario Vignali, University of Pittsburgh School of Medicine) were used. MC38 cells were grown in DMEM supplemented with 10% FBS, 100 U penicillin, and 100 μg streptomycin in a 37°C incubator with 5% CO_2_. Cells were harvested at passage 2-3 after thaw, and 2.5×10^5^ tumor cells were injected subcutaneously into the flank of mice anesthetized with 2.5% 2,2,2,-Tribromoethanol (Avertin). Tumors were measured every 2-3 days using calipers, and mice were sacrificed when tumors reached 2 cm^3^ volume, ulceration, or a body condition of >2 in accordance with IACUC guidelines. Tumor volume was determined using the formula for the volume of an ellipsoid, ½ × D × d^2^, where “D” is the major axis of the tumor and “d” is the minor axis. Tumors were harvested from mice at days 18-23 after implantation for single cell RNA sequencing experiments and flow validation experiments as indicated in the Figure Legends.

#### Clinical Samples

Studies of patients with melanoma were approved by the UCSF Committee on Human Research (CC138510) and by the Institutional Review Board of UCSF under protocol 13-12246. All patients provided written, informed consent prior to biopsy and/or blood collection.

### METHOD DETAILS

#### Lymphocyte isolation from mouse tissues

Peripheral blood was collected from mice using the retroorbital bleeding route, and blood was collected into 4% sodium citrate (Sigma) to prevent clotting. RPMI+10% FBS was added to dilute out the anti-coagulant, and then white blood cells were separated from red blood cells using centrifugation through histopaque-1083 (Sigma). The white blood cell layer at the interface between the histopaque and remaining media was subsequently washed and subjected to staining for flow cytometry analysis or sorting for single cell RNA sequencing.

Tumors were dissected and mechanically disaggregated. For flow cytometry validations, a GentleMACS (Miltenyi) was used for disaggregation, whereas for single cell RNA sequencing vertical scissors used to mince the tumors instead of the GentleMACS. The dissociated tissue was digested with Collagenase Type I (400 U/ml; Worthington Biochemical) for 20-30 minutes at 37°C. Samples were then passed through a 70 μm filter, and lymphocytes were enriched using centrifugation through a Percoll gradient (40% and 70%). The enriched lymphocyte layer at the 40%/70% interface was subsequently washed and stained for flow cytometry or sorted for single cell RNA sequencing.

#### Flow cytometry and sorting of mouse samples

Single cell suspensions were generated as described above. Suspensions were labeled with LIVE/DEAD Fixable Near-IR Cell Stain in PBS (Thermo Fisher Scientific) to exclude dead cells from downstream analyses. Cells were pre-incubated with TruStain Fc Receptor Block (anti-mouse CD16/CD32, clone 93, BioLegend), then labeled with extracellular antibodies including: CD3 (clone 145-2C11) and CD8a (clone 53-6.7) (from BD); CD11a (clone M17/4) (from Thermo Fisher Scientific); CCR2 and NKG2I (R&D Systems); Lag3 (clone C9B7W) (from Bio-Rad); and CD45.2 (clone 104), PD-1 (clone RMPI-30), CX3CR1 (clone SA011F11), CD62L (MEL-14), CD44 (IM7), CCR5 (clone HM-CCR5), CXCR6 (clone SA051D1), CD49D (clone R1-2), CD18 (clone M18/2), CD29 (clone HMβ 1-1), CD48 (clone HM48-1), CD94 (clone 18d3), NKG2D (clone CX5 or C7), CD39 (clone Duha59), NKG2A (clone 16A11), NK1.1 (clone PK136), Tim-3 (clone RMT3-23), CD160 (clone 7H1), Slamf7 (clone 4G2), TIGIT (clone IG9), and NRP1 (clone 3E12) (from BioLegend). Flow cytometry labeling (without inclusion of Feature Barcoding antibodies from BioLegend) was performed in PBS supplemented with 2% FBS. For CITE-seq validation experiments, cells were labeled with TotalSeqC antibodies against CD39 (TotalSeq C0834, clone Duha59) and CX3CR1 (TotalSeq C0563, clone SA011F11) as directly conjugated antibodies, and NKG2D as a biotin/streptavidin reaction (NKG2D-biotin clone C7 paired with TotalSeq C0971-Steptavidin) (from BioLegend). Labeling with Feature Barcoding antibodies was performed in PBS supplemented with 2% BSA and 0.01% Tween. Samples were acquired on a FACSymphony (BD Biosciences) and analyzed with Flow Jo software (BD Biosciences). Flow cytometry-based sorting for single cell RNA seq was performed using a FACSAria (BD Biosciences). Blood samples were sorted based on live, CD45.2^+^, CD3^+^, CD8α^+^, CD44^high^. Tumor samples were sorted based on live, CD45.2^+^, CD3^+^, CD8a^+^.

#### Single cell RNA sequencing of mouse samples

Gene expression and TCR libraries for mouse samples were generated using the Chromium Single Cell 5’ Library and V(D)J Reagent Kit (10X Genomics) according to the manufacturer’s recommendations. For samples requiring Feature Barcoding libraries to detect TotalSeqC antibodies (from BioLegend), the Chromium Single Cell 5’ Feature Barcode Library Kit (10X Genomics) was used according to the manufacturer’s recommendations. Following sorting as described above, approximately 10,000 cells per sample were loaded into each channel of the Chromium Chip, and recommendations were followed assuming targeted cell recovery of 2,001-6,000 cells. Libraries were sequenced on a NextSeq sequencer (Illumina) by the Dana-Farber Cancer Institute Sequencing Core. Gene expression libraries and Feature Barcoding libraries were sequenced using the 26 × 8 × 91 bp parameters recommended by 10X Genomics. TCR libraries were sequenced using the 150 × 8 × 150 bp parameters recommended by 10X Genomics. Based on approximate cell numbers expected, we sequenced a minimum of 20,000 reads per cell for gene expression libraries and 5,000 reads per cell for TCR and Feature Barcoding libraries.

#### Lymphocyte isolation from human tissue samples

Human melanoma tumor samples were mechanically dissociated and enzymatically digested overnight for 12-14 hours. Following fine mincing with scissors, samples were digested in RPMI media (Gibco) containing 250 U/mL Type IV collagenase (4188; Worthington Biochemical Corp.), 20 μg/mL DNAse (SDN25-1G; Sigma-Aldrich), 10%FBS (Alphabioregen), 1% HEPES (Gibco), 1% penicillin/streptomycin (Gibco), and 2mM glutamine (GLUTAmax, Gibco) at 37°C in a tissue culture incubator with 5% CO_2_. Following overnight incubation, digestion was quenched with excess media, and samples were transferred to 50mL conical tubes and briefly shaken, and then were filtered through a 100 μm sieve. Samples were pelleted and washed in media before downstream applications.

#### Lymphocyte isolation from human blood samples

Blood from patients with melanoma was collected in heparinized or EDTA tubes and was diluted with an equal volume of PBS before being layered over a Ficoll Paque PLUS gradient (GE Healthcare) in 50 mL conical tubes that were centrifuged at room temperature for 15 minutes at 932 g. Cells were isolated from the Ficoll/PBS interface and were washed at least twice in PBS/2%FBS before downstream applications.

#### Flow cytometry and sorting of human samples

Melanoma tumors (primary tumors or metastases as indicated in Fig. S4B and Table S7) or blood were stained in PBS with Tonbo Ghost Dye Violet 510, anti-CD45 (Clone H130), anti-CD3 (Clone SK7), anti-CD4 (Clone SK3), anti-CD8 (clone SK1). Some samples were additionally stained with anti-PD-1 (Clone EH12.2H7), anti-CD25 (Clone M-A251), anti-CD27 (Clone LG.7F9), anti-CD127 (Clone HIL-7R-M21). CD8 T cells were sort-purified as singlet, live, CD45^+^, CD3^+^, CD4^−^, CD8^+^ events on an Aria 2 or Aria 3u (BD) in the UCSF Parnassus Flow Cytometry Core. In some cases the total CD3^+^ T cell population was sort-purified as singlet, live, CD45^+^, CD3^+^ events, and CD8^+^ T cells were identified bioinformatically. Cells were counted post-sort on a hematocytometer and resuspended to target ~1000 cells/μL in media with 10% FBS for single-cell RNA Sequencing.

#### Single cell RNA sequencing of human samples

Following sorting, cells were prepared for single-cell RNA Sequencing using the 10x Chromium Platform (10x Genomics) by the Institute for Human Genetics at UCSF. Cells were processed following the recommended protocol with the Chromium Single Cell 5' Library Construction kit and Chromium Single Cell V(D)J Enrichment Kit (Human T Cell) (Single Cell 5' PE Chemistry). Libraries were run on a HiSeq 4000. FASTQ files were generated and analyzed with Cell Ranger (v3.0.2) by the UCSF 10x Genomics Core using the GRCh38 human reference genome for alignment.

#### Demultiplexing and Read Processing

Raw reads were processed using Cell Ranger v3.0.2 to generate raw counts matrices of gene expression and csv files corresponding to TCR clonality. The GRCh38 human reference genome was used for alignment of human samples, and the mm10 mouse reference genome was used for alignment of mouse samples. Aether version 1.0 (Luber et al., 2018) was used to process certain resource heavy jobs on compute instances rented from Amazon Web Services.

#### Computational Processing of Gene Expression data

All analyses were conducted using R version 3.6.1 and Seurat version 3 with additional utilization of the dplyr, data.table, ggplot2, cowplot, viridis, gridExtra, RColorBrewer, ggpubr, ggrepel, gtools, DescTools, doParallel, doSNOW, and tibble packages. Seurat objects were created with the min.cells parameter set to 3 and the min.features parameter set to 400. Filtering cells based on expression of housekeeping genes was conducted using the human and mouse (where appropriate) gene lists maintained by the Seurat developers (available on the Satija lab website), with cells passing the filtering criteria if they had expression greater than 0 for more than half of the genes in the list. Subsequently, the MitoCarta database from the Broad institute was utilized to filter out cells based on expression of mitochondrial genes (Calvo et al., 2016). Cells were filtered out if they expressed more than 500 of the 1158 mitochondrial genes in human, or if the number of mitochondrial genes expressed was higher than 2 standard deviations from the mean in mouse.

Data were normalized using the default Seurat function (generating log-transformed transcripts-per-10K read measurements) followed by scaling, and variable genes were found using “ExpMean” for the mean.function parameter and “LogVMR” for the dispersion.function parameter. The RunPCA function was run utilizing 50 principal components and then the FindNeighbors function was run using 30 dimensions. Subsequently, the FindClusters function was run with a resolution aiming to generate 5-7 biologically meaningful clusters per sample. To filter for CD8^+^ T cells in humans, clusters were kept if (1) the proportion of cells in the cluster with at least two genes out of *CD3E*, *CD3D*, or *CD3G* being expressed is greater than 30% and either (2) *CD8B* is expressed in more than 30% of cells in the cluster, *CD8A* is expressed in more than 30% of cells in the cluster, *FOXP3* is expressed in less than 5% of cells in the cluster, and *CD4* is expressed in less than 5% of cells in the cluster or (3) *MKI67* is expressed in greater than 70% of cells in the cluster and either *CD8A* or *CD8B* is expressed in greater than 20% of the cells in the cluster. This last criterion is to account for proliferating clusters. In mice, which had less contamination from non-CD8+ T cells due to prior sorting, clusters were kept if more than 30% of cells in the cluster expressed any of *Cd3e*, *Cd3d*, or *Cd3g* and: 1) if more than 30% of cells in the cluster expressed *Cd3e* and *Cd8a* while having less than 5% of the cells express *Foxp3*. When applicable, samples were integrated using the SCTransform method (Hafemeister and Satija, 2019). Upon obtaining transcriptional clusters in the integrated datasets, up regulated genes associated with each cluster were determined via the Wilcox Rank Sum test implemented in the FindAllMarkers function in Seurat.

Cells were classified as positive for the PD-1 transcript (*Pdcd1* in mice, *PDCD1* in humans) if they had any number of reads above zero. To classify mouse cells as positive for the *Klrk1* (encoding NKG2D)*, Entpd1* (encoding CD39), and/or *Cx3cr1* (encoding CX3CR1), a more stringent cut off was used for a cell to qualify as positive, determined by COMET (Delaney et al., 2019). The COMET-determined thresholds for all markers in the COMET database are provided for the mouse dataset in Table S4, and the human dataset in Table S10, and can be found in the column titled “cutoff_val”.

For enrichment analysis tests (Fig. 1E), all genes were ranked by their p-value and by their fold change, and then the two ranking values were aggregated to create a single ranking by taking the mean of the p-value and fold-change rankings. We then search for significant associations with gene signatures by using the ranked list in the preranked analysis of GSEA. Default settings were used except: permutations was set to 100, the enrichment statistic set to ‘classic’, and the max size set to 2500. The signature sets used were all GO terms, Kegg and Reactome pathways, and immune signatures from MSigDB (groups c2, c5, and c7). Gene signatures derived from the literature were also analyzed as cited in the figures and manuscript.

In order to perform the clonal-corrected DE gene analysis comparing TM cells in the blood to blood-matching cells in the tumor, the non-normalized integrated mouse blood object was subset to keep only TM cells, which were then collapsed into their clones such that for each gene, the counts for all the cells in a clone were summed together. This was done for the integrated tumor as well, but with blood-matching cells. The tumor and blood-derived datasets were then merged to a single object, and the edgeR package was used to call differential expression. Genes were considered if they expressed at least 1 count per million, and then counts were normalized using the trimmed mean of M-values. Taking into account the paired nature of matching clones in blood and tumor, genes were fit to a generalized linear model using the ‘glmFit’ function, and likelihood ratio tests were conducted to detect DE genes between blood and tumor with the ‘glmLRT’ function.

#### Single-cell TCR and clonal analysis

Cells for which at least one alpha and one beta chain were annotated in the TCR data were determined as “tumor/blood-matching” or “tumor/blood-non-matching” based on whether there was a cell in the paired tissue data that had the exact same alpha and beta chain composition as the given cell. Only cells that had at least one alpha chain and one beta chain annotated were included in all of the analyses and visualizations comparing “matching” to “non-matching” cells. Two cells were assigned to be in the same clone if they had the both the exact same alpha and beta chains assigned based on the amino acid sequence. If cells had more than one alpha and beta chain, they were considered matching if all of the alpha and beta chains detected were shared. This strict definition was used to ensure each pair of cells within the same clone has complete similarity of the TCR chains detected, and hence is with high probability derived from the same T cell clone.

To define tumor-matching status by clustering of TCRs, two algorithms were employed: GLIPH2 (Huang et al., 2020) and iSMART (Zhang et al., 2020). For each patient, the joint collection of blood and tumor CD8^+^ TCRs were submitted to each algorithm individually for clustering on default parameters. All resultant clusters that included at least one TCR found in the tumor sample were considered to represent reactivity to a tumor antigen, and therefore all blood CD8^+^ T cells with TCRs belonging to these clusters were considered TM cells. In general, the results of GLIPH2 and iSMART were concordant with 4,557 cell TM labels in agreement and 74 in disagreement. To buffer this analysis against variation in algorithm and parameter choices, we disregarded the 74 cells for which the two algorithms gave conflicting results (<1.6% of cells).

#### Functional Annotations of Seurat Clusters

Functional annotations for Seurat clusters were manually curated using a combination of up regulated genes for each cluster and visual inspection of key markers using UMAP visualization. Key markers used for aiding in annotation included: *Sell, Tcf7, Lef1, Ccr7, Il7r, S1pr1, Klf2, Cxcr3, Klrg1, Cx3cr1, S1pr5, Tnf, Ifng, Il2ra, Gzmb, Prf1, Mki67, Slamf6, Pdcd1, Lag3, Tigit, Cd160, Havcr2, Ctla4, Bst2, Irf1, Irf2, Irf7, Mx1, Ccr6, Rorc, Cxcr6, Itgae, cd69, Tbx21,* and *Eomes.* Transcriptional signatures in blood consistent with naïve, central memory, effector, and effector memory cells, and signatures in tumor consistent with diverse exhausted subsets, effector-like, resident memory-like, naïve/central-memory like, IFN-stimulated, and cycling populations, as previously reported (Best et al., 2013; Guo et al., 2018; He et al., 2016; Im et al., 2016; Kakaradov et al., 2017; Kurtulus et al., 2019; Miller et al., 2019; Milner et al., 2017; Sade-Feldman et al., 2018; Siddiqui et al., 2019; Tirosh et al., 2016; van der Leun et al., 2020; Yost et al., 2019).

Clusters that expressed high levels of *Sell, Tcf7, Ccr7, Il7r, S1pr1,* and *Klf2* and lower levels of *Klrg1, Cx3cr1, S1pr5, Tnf, Ifng, Gzmb, Prf1, Mki67,* and the inhibitory receptors (e.g. *Pdcd1, Havcr2, Ctla4*, etc.) were considered naïve and/or central-memory like. Clusters that expressed high levels of *Klrg1, Cx3cr1, S1pr5, Tnf, Ifng, Gzmb,* and *Prf1* and low levels of *Sell, Tcf7, Lef1, Ccr7,* and *Il7r* were considered effector and/or effector-memory like. Exhausted subsets were classified as those expressing multiple inhibitory receptors (*Pdcd1, Havcr2, Lag3, Tigit*, etc.), low levels of naïve and/or central memory-like markers, and generally lower levels of some effector molecules such as *Klrg1* and *Cx3cr1*. The exhausted populations were further subdivided into progenitor-like (based on expression of *Tcf7* and lower levels of *Havcr2*), intermediate-like (based on low levels of *Tcf7* and *Havcr2* and expression of other IRs including *Pdcd1, Ctla4, Lag3, CD160*, etc.), and terminal-like (based on high levels of multiple inhibitory receptors including *Pdcd1, Havcr2, Ctla4, Lag3, Cd160,* etc.). An interferon-stimulated cluster was defined based on over representation of IFN responsive genes in the up regulated gene list, including *Bst2, Irf1, Irf2, Irf7, Stat1, Stat2,* and *Mx1.* Clusters containing cells that were undergoing cell cycle were identified based on over representation of cell cycle genes (including *Mki67* and several *Kif, Cdk, and Cdc* genes). Lastly, resident memory-like populations were identified based on expression of *Itgae*, *Itga1,* and *Cxcr6*.

#### Transcriptional Signature Analysis

We computed the extent to which gene signatures are expressed in cells by using Scanpy’s ‘score_genes’ function on the centered and scaled gene count data objects (Wolf et al., 2018). Because gene signature computation is relative (following centering and scaling of the gene expression data), data of all cells compared was merged prior to the centering and scaling procedure. Violin plots were generated with the ‘seaborn’ package in Python.

To create the plot shown in Fig. S3C, all cells were merged into a single data object and normalized to 10,000 counts per cell. Only cells with at least one alpha and one beta chain were included. Then each count was logarithmized according to log(1+X), where X is the gene count, and each gene was standardized to unit variance and zero mean. Given a signature, a score was calculated for each cell with Scanpy’s ‘score_genes’ function. The average of the cell scores was calculated for each sample.

#### Machine Learning

Classification of tumor-matching cells in mouse (Fig. 2A, 2B) was conducted using L2 regularized logistic regression using the Scikit-learn package in python version 2.8 (Pedregosa F., 2011). Plots were generated using matplotlib. For the Logistic regression, the liblinear solver was used with a l2 penalty and C parameter set to .02.

#### Calculation of AUC

For each gene in each patient, the area under the curve in distinguishing tumor-matching cells from non-tumor-matching cells was computed with the AUC() function of the R DescTools package. To construct an input for the AUC() function, we calculated a vector of (1-Specificity) values and a vector of corresponding Sensitivity values from 39 potential expression level thresholds for dividing the two populations. For each gene in each patient, the 39 thresholds were every 5^th^ percentile expression of the gene (21 values including 0^th^ percentile and 100^th^ percentile) combined with 18 evenly spaced expression values between the minimum and maximum, to account for heavily skewed distributions in which useful thresholds may lie above the 95^th^ or below the 5^th^ percentile. To these input vectors we added (0,0) and (1,1), representing the trivial options of selecting none and all of the cells as TM, respectively.

#### Similarity of TM Component Across Patients

Similarity between samples in terms of transcripts’ power in distinguishing the TM component from the non-TM component was computed in Fig. 5C via pairwise correlation of gene transcript AUC values for selecting the TM component. The AUC for each gene transcript in each patient was calculated as described above, and all pairwise combinations between patient samples were plotted for each gene, resulting in 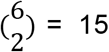 points per gene. Negation markers are represented by AUC values < 0.5 when selecting the TM component. To mitigate the x axis being arbitrarily biased toward the patients appearing first in the data, x and y coordinates were switched for each point with a probability of 0.5 and a random seed set to 27 in R. With the function stat_cor() from R package ggpubr, Pearson correlation statistics were computed for the resultant x and y values, stratified by if each sample pair was within the same patient or different patients. The plot is restricted to transcripts measured in all six patient samples.

#### COMET

COMET (Delaney et al., 2019) runs were conducted with version 0.1.12 the default X parameter (0.15) and with the L parameter set to the minimum of (1) 10*K and (2) 0.35*N, where K is the number of tumor matching cells and N is the total number of cells with at least one alpha and one beta chain annotated, to account for our willingness to allow for greater levels of contamination in the identified tumor-matching samples than allowed by default.

The full list of unranked markers from COMET are provided in Table S4 for mouse, and Table S10 for human. In these files, the COMET-determined threshold value (column labeled “cutoff_val”) is indicated for each marker when used as a positive marker or a negation marker. Negation markers are labeled as “marker_negation”, whereas positive markers are simply listed at “marker”. For positive markers, a cell is predicted to be TM if its gene expression is above the threshold. For negation markers, a cell is predicted to be TM if its gene expression is lower than the threshold. In COMET’s original output, negation markers are multiplied by (−1). We therefore took the absolute value of all reported thresholds in the output tables to increase clarity. Since “negation” does not necessarily equate to no expression, throughout the text cells deemed positive for a “negation marker” are referred to as “marker low” instead of “marker negative.”

#### Ranking singleton human markers

Leading candidate markers for follow-up analysis from the human samples were determined by the number of patient samples in which a given marker reached significance in COMET (q <0.05). From the input list (Chihara et al., 2018), we removed *CD8A* because this is a lineage defining marker and therefore not ideal for separating the TM and non-TM components, along with cytokines *CCL4*, *CCL5*, and *MIF* in order to strictly consider surface-expressed markers. 15 markers derived from this filtered list had q < 0.05 in the majority of patient samples (4 of 6) and were considered for follow-up analysis. These 15 candidate markers were ranked in order of descending AUC in distinguishing TM cells from non-TM cells for each patient, and ranks were averaged across patients to reach a summary continuous ranking for each of the 15 markers (Table S12). The top four on this list were negation markers for *LTB, CCR7, GYPC,* and *FLT3LG*.

#### Empirical p-values and confidence intervals for gate performance

Confidence intervals for gate AUC, sensitivity, and specificity were determined by 10,000 iterations of random bootstrap resampling with replacement in the pooled CD8^+^ blood cell population and separately with respect to each individual patient blood sample. The 95% confidence intervals given go from the 2.5^th^ percentile to the 97.5^th^ percentile of 10,000 bootstrapped recalculations of AUC, sensitivity, and specificity. A null distribution for each gate was generated iteratively through each resample by permuting the “tumor-matching” labels and calculating the AUC, sensitivity, and specificity from the resultant datasets. These distributions represented the null hypothesis that each given marker was sorting TM cells from non-TM cells by chance alone. Reported empirical p-values < 0.0001 reflect the observation that the point estimate for the marker’s AUC, sensitivity, or specificity was never observed in the 10,000 iterations of the null.

#### Ranking of combinatorial marker gates

All possible one, two, three, and four gene logical gates were enumerated from the four top-ranking markers in the patient samples (*CCR7*^low^, *FLT3LG* ^low^, *GYPC*^low^, and *LTB* ^low^) and evaluated for their sensitivity and specificity in isolating the TM component in each patient at a universal threshold of 0.001 UMIs. The optimal threshold to discriminate these two populations must be calibrated to the distribution of read counts as well as the target sensitivity and specificity. We used COMET to determine this optimal threshold for each marker in each patient sample (Table S14), and chose a universal threshold of 0.001 following manual inspection (COMET-derived thresholds averaged across patient samples were 0.001, 0.001, 0.832, and 0.332 for *CCR7, FLT3LG, GYPC,* and *LTB,* respectively, and any threshold between 0 and 1 is functionally equivalent when applied to count data). To identify the best-performing combinatorial gate, we computed a penalty for each gate based on both its distance from perfect sensitivity and specificity as well as its balance between the two metrics. To calculate this penalty, we first computed the Euclidean distance from perfect sensitivity and specificity (corresponding to (0,1) on ROC curve plots) to the point on the plot representing that gate’s sensitivity and specificity in the pool of CD8^+^ cells across all patient blood samples. To this Euclidean distance, we added the difference between the gate’s sensitivity and specificity in the pool of CD8^+^ T cells across all patient blood samples in order to promote the selection of the most balanced gate. This process identified [*(CCR7^low^ and FLT3LG^low^ and GYPC^low^) or LTB^low^*] as the best-performing and most balanced gate (lowest penalty).

### QUANTIFICATION AND STATISTICAL ANALYSIS

#### Flow cytometry validations in mouse

Statistical analyses for flow cytometry data were performed with Prism software (GraphPad), and p-values of less than 0.05 were considered statistically significant. Multiple t tests were performed using the Holm-Sidak method with alpha = 0.05. Each row was analyzed individually, without assuming a consistent SD. Asterisks indicating significance in the Figures correspond to p<0.05 (*), p<0.01 (**), and p<0.001 (***). Statistical tests used for computational analyses are indicated in the corresponding Figure Legends and Methods sections. Exact p values for significant comparisons are indicated in the Figure Legends and Supplemental Tables.

### RESOURCE AVAILABILITY

Further information and requests for resources and reagents should be directed to and will be fulfilled by the corresponding authors. This study did not generate new unique reagents. The gene expression scRNA seq data for patients K409 and K411 (initial blood/tumor pair) and the TCR data (for K409 tumor) can be found on GEO with accession number GSE148190 (Mahuron et al., 2020). The scRNA-seq data generated during this study will be made available upon publication.

### DESCRIPTION OF SUPPLEMENTAL MATERIAL

This work contains five Supplemental Figures (Fig. S1-S5) and fifteen supplemental excel tables to support the data in the Main Figures. Figure S1 is associated with Figure 1, and provides additional details characterizing the single-cell RNA seq discovery dataset in mice. Figure S2 is associated with Figure 2, and provides additional information about the COMET output, flow cytometry validations in mice, and combinations of NKG2D, CX3CR1, and CD39 in our flow cytometry data and CITE seq data. Figure S3 is associated with Figure 3, and provides additional comparisons between matching clones in blood and tumor in replicate mice. Figure S4 is associated with Figures 4 and 6, and provides supporting information regarding the single-cell RNA seq in the melanoma patients. Figure S5 is associated with Figure 4–6, and details results from alternative methods to identify matching clones based on the TCR in melanoma patients. A list detailing the fifteen supplemental excel tables is provided.

## Supplemental Tables

**Table S1**: Up regulated genes for each Seurat cluster in mouse integrated blood and MC38 tumor samples.

**Table S2:** Up regulated genes for tumor matching and non-matching CD8^+^ T cells in the peripheral blood of mice with MC38 tumors

**Table S3**: Full list of pathways and signatures enriched in tumor matching and in non-matching CD8^+^ T cells from the peripheral blood of mice with MC38 tumors

**Table S4:** Significance measures calculated with COMET in sorting tumor matching from non-matching CD8^+^ T cells in the blood of mice with MC38 tumors

**Table S5:** Sensitivity and specificity of all possible gates made from combinations of NKG2D, CD39, and CX3CR1, measured by CITE seq in mice

**Table S6:** Differentially expressed genes between tumor-matching cells in blood and blood-matching cells in tumor

**Table S7:** Clinical parameters for patient samples

**Table S8**: Up regulated genes for each Seurat cluster in human integrated blood and initial tumor samples

**Table S9:** Transcript AUC performance, delineated by melanoma patient sample

**Table S10:** Significance measures calculated with COMET for all transcripts in sorting tumor matching from non-matching CD8^+^ T cells in the blood of melanoma patients

**Table S11:** Similarities and differences in COMET-identified markers to identify TM cells in mice with MC38 tumors compared to human melanoma patients

**Table S12:** The 15 transcripts that were significant in at least 4 patient samples, ordered by average ranking of their AUC

**Table S13:** Empirical significance values and 95% confidence intervals for the AUC, sensitivity, and specificity of featured gates in each patient sample

**Table S14:** Sensitivity and specificity values for all possible transcriptional marker combinations of *CCR7*^low^, *GYPC*^low^, *FLT3LG*^low^, and *LTB*^low^, delineated by patient sample

**Table S15**: Empirical significance values and 95% confidence intervals for the AUC, sensitivity, and specificity of featured gates in each patient sample using the intersection of GLIPH2 and iSMART

