## Supplemental Figures for "Single-cell analyses identify circulating anti-tumor CD8 T cells and markers for their enrichment"

Figure S1

a

| Blood Sample | # of cells (GEX) | # of cells (GEX+TCR) | # Matching cells | % Matching cells | % *Pdcd1*+ TM cells |
|---|---|---|---|---|---|
| M1 | 3124 | 1815 | 183 | 10.08 | 11.48 |
| M2 | 3647 | 1803 | 22 | 1.22 | 9.09 |
| M3 | 3509 | 2088 | 28 | 1.34 | 10.71 |

b

| MC38 tumor Sample | # of cells (GEX) | # of cells (GEX+TCR) | # Matching cells | % Matching cells | % *Pdcd1*+ TM cells |
|---|---|---|---|---|---|
| M1 | 3135 | 2185 | 752 | 34.42 | 74.60 |
| M2 | 3167 | 2407 | 1144 | 47.53 | 94.49 |
| M3 | 2148 | 1779 | 239 | 13.43 | 85.77 |

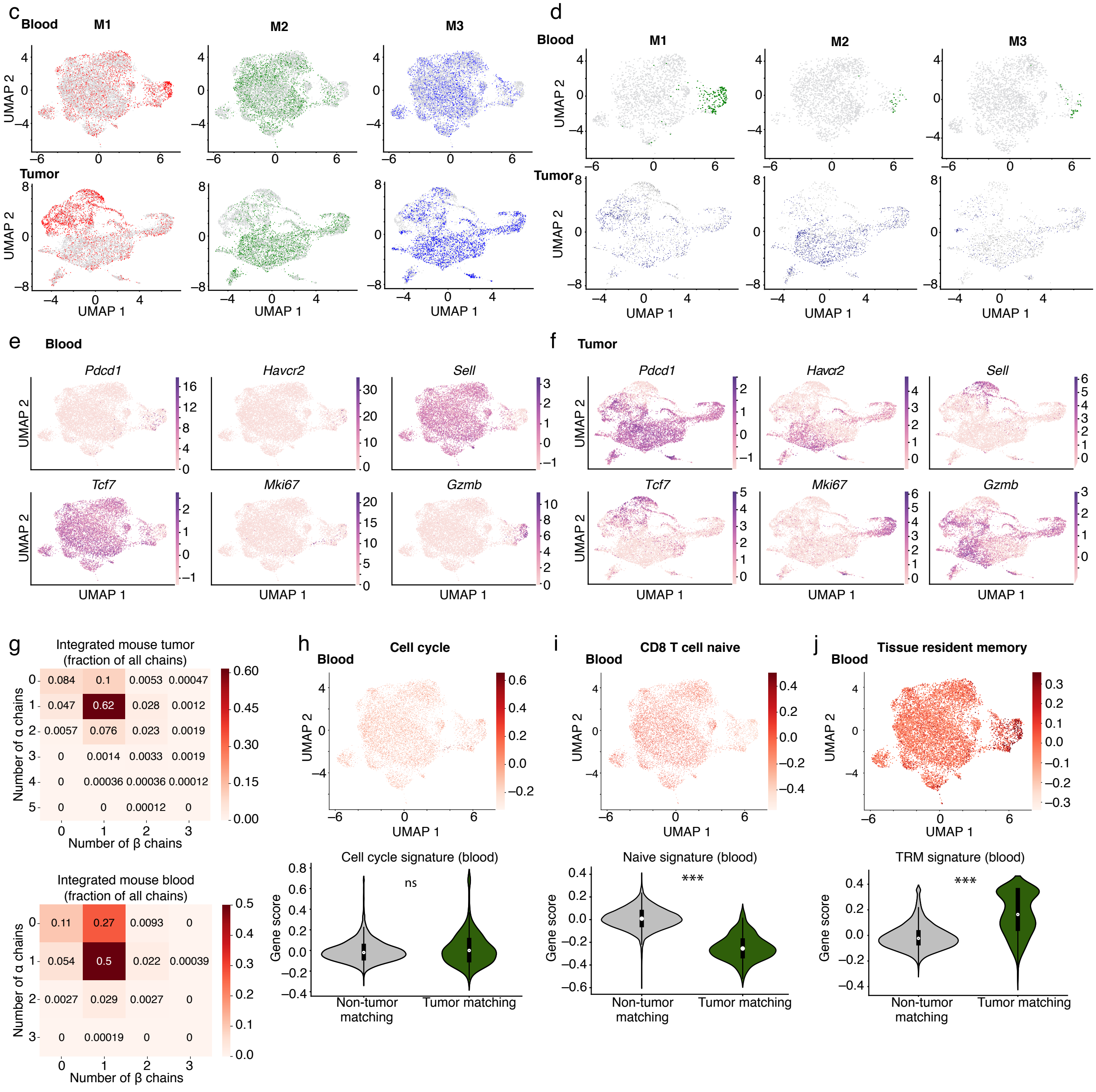

**Figure S1: Transcriptional landscape of CD8$^+$ T cells in paired peripheral blood and MC38 tumors in mice.** (a-b) Tables indicating details about each mouse in the discovery 10X cohort (Mouse 1 (M1), Mouse 2 (M2), and Mouse 3 (M3)), including the number of cells recovered that had gene expression (GEX) data, GEX and TCR data, number of matching cells, percentage matching cells of the total sorted population, and the frequency *of Pdcd1*$^+$ TM cells in (a) peripheral blood and (b) MC38 tumors. (c) UMAP of the integrated blood samples (Top) and MC38 tumor samples (Bottom) showing the distribution of each mouse in the integrated dataset (datasets combined from M1, M2, and M3). Cells from each mouse shown in color (M1 = red, M2 = green, M3 = blue), and the cells from the other two mice are shown in grey for each plot. (d) UMAP of the integrated blood samples (Top) and MC38 tumor samples (Bottom) showing the distribution of clones shared between tissues (TM cells in blood, and blood-matching cells in tumor). Only TM cells (green), blood-matching cells (navy blue), and non-matching cells (grey) from each individual mouse are shown, and the cells from the other two mice in the integrated object are excluded. (e-f) UMAPs showing distribution of expression of select transcripts in the integrated blood (e) and MC38 tumor (f) samples. Genes include *Pdcd1* (encoding PD-1)*, Havcr2* (encoding Tim-3)*, Sell* (encoding CD62L)*, Tcf7* (encoding TCF-1)*, Mki67* (encoding Ki-67)*,* and *Gzmb* (encoding granzyme B)*.* (g) Heat map showing the fraction of cells in the integrated MC38 tumor (Top) and blood (Bottom) datasets with the indicated number of TCR alpha and beta chains detected. (h-j) (Top) UMAP of integrated blood samples showing expression of (h) a cell cycle signature (Kowalczyk et al., 2015) (p=0.24), (i) a CD8$^+$ naïve T cell signature (Kaech et al., 2002) ($p=4.6x10^{-125}$), and (j) a tissue resident memory signature (Beura et al., 2018) ($p=2.6x10^{-59}$). Violin plots quantifying the expression of each signature in (h-j) in TM compared to non-TM cells in the blood (Bottom). Significance determined using Wilcoxon rank-sum tests.

Figure S2

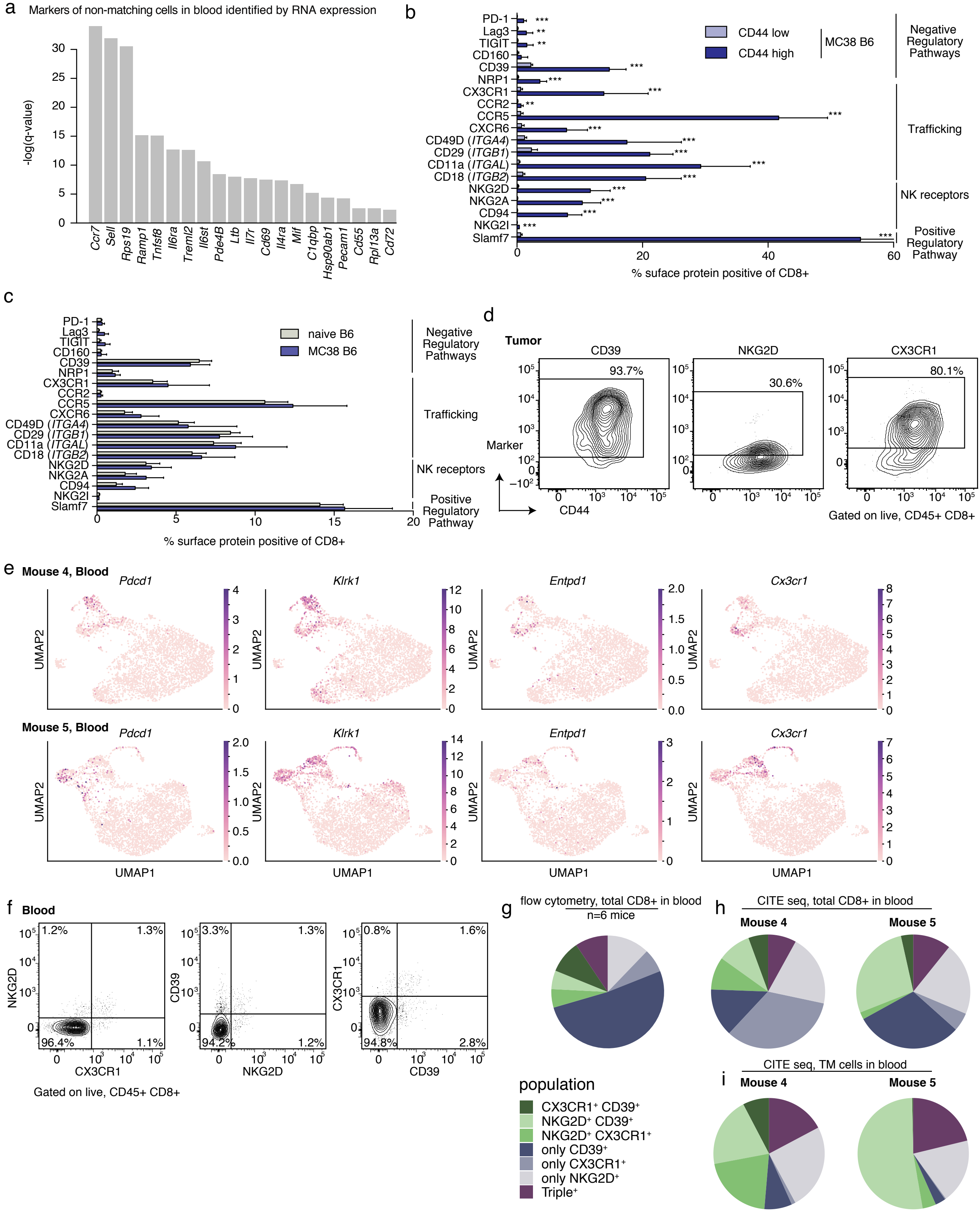
a
Markers of non-matching cells in blood identified by RNA expression
-log(q-value)
Ccr7
Sell
Rps19
Ramp1
Tnfsf8
Il6ra
Treml2
Il6st
Pde4B
Ltb
Il7r
Cd69
Il4ra
Mif
C1qbp
Hsp90ab1
Pecam1
Cd55
Rpl13a
Cd72
b
PD-1
Lag3
TIGIT
CD160
CD39
NRP1
CX3CR1
CCR2
CCR5
CXCR6
CD49D (ITGA4)
CD29 (ITGB1)
CD11a (ITGAL)
CD18 (ITGB2)
NKG2D
NKG2A
CD94
NKG2I
Slamf7
CD44 low
CD44 high
MC38 B6
Negative Regulatory Pathways
Trafficking
NK receptors
Positive Regulatory Pathway
% suface protein positive of CD8+
c
naive B6
MC38 B6
Negative Regulatory Pathways
Trafficking
NK receptors
Positive Regulatory Pathway
% surface protein positive of CD8+
d
Tumor
CD39
NKG2D
CX3CR1
93.7%
30.6%
80.1%
Marker
CD44
Gated on live, CD45+ CD8+
e
Mouse 4, Blood
Mouse 5, Blood
Pdcd1
Klrk1
Entpd1
Cx3cr1
UMAP2
UMAP1
f
Blood
1.2%
1.3%
96.4%
1.1%
NKG2D
CX3CR1
3.3%
1.3%
94.2%
1.2%
CD39
NKG2D
0.8%
1.6%
94.8%
2.8%
CX3CR1
CD39
Gated on live, CD45+ CD8+
g
flow cytometry, total CD8+ in blood
n=6 mice
h
CITE seq, total CD8+ in blood
Mouse 4
Mouse 5
i
CITE seq, TM cells in blood
Mouse 4
Mouse 5
population
CX3CR1⁺ CD39⁺
NKG2D⁺ CD39⁺
NKG2D⁺ CX3CR1⁺
only CD39⁺
only CX3CR1⁺
only NKG2D⁺
Triple⁺

**Figure S2: Identification and validation of markers to identify tumor-matching T cells in blood.** (a) Top surface markers for identifying non-TM cells from TM cells in the blood based on COMET (Delaney et al., 2019) analysis. Significance determined using an XL-minimal hypergeometric test with multiple hypothesis test corrections. (b) Quantification of the frequency of bulk $CD8^{+}$ T cells in the peripheral blood of mice with MC38 tumors at d21 post implantation (n=9 mice) that express the indicated proteins using FACS. Cells are gated on singlets, live/dead$^{-}$, $CD45^{+}$, $CD8\alpha^{+}$, and are further gated based on CD44 expression to compare $CD44^{low}$ and $CD44^{high}$ cells. (c) Comparison of bulk $CD8^{+}$ T cells (gated on singlets, live/dead$^{-}$, $CD45^{+}$, $CD8\alpha^{+}$) from the peripheral blood of mice with MC38 tumors at day 21 post implantation (n=9 mice) to naïve B6 mice (n=4 mice). For (b-c), data are representative of 2-4 independent experiments depending on the marker with n=3-4 naïve mice and n=5-9 mice with MC38 tumors (d19-22). Bars show the mean, and error bars represent SD. Significance determined using multiple t tests using the Holm-Sidak method, with alpha = 0.05. Each row was analyzed individually, without assuming a consistent SD. Reported are the adjusted p values considering multiple tests. Significant comparisons in (b) indicated with asterisks and include PD-1 $p=2.6957 \times 10^{-5}$, Lag-3 p=0.0012, TIGIT p= p=0.0012, CD39 $p=3.7639 \times 10^{-9}$, NRP1 $p=5.1172 \times 10^{-7}$, CX3CR1 p=0.0002, CCR2 p=0.0012, CCR5 $p=6.4414 \times 10^{-10}$, CXCR6 $p=6.1903 \times 10^{-5}$, CD49 p=0.0002, CD29 $p=1.6745 \times 10^{-9}$, CD11a $p=1.1636 \times 10^{-7}$, CD18 $p=1.9906 \times 10^{-7}$, NKG2D $p=1.1863 \times 10^{-7}$, NKG2A $p=1.8567 \times 10^{-7}$, CD94 $p=1.8567 \times 10^{-7}$, NKG2I $p=2.6625 \times 10^{-6}$, Slamf7 $p=5.06 \times 10^{-13}$. In (b), CD160 expression between CD44 high and CD44 low was not significant. In (c), there were no significant differences between naïve B6 and B6 mice with MC38 tumors. (d) Representative FACS contour plots showing NKG2D, CD39, and CX3CR1 expression (Y axis) as indicated above each plot, and CD44 (X axis) on $CD8^{+}$ T cells in the MC38 tumor of mice in Fig. 2f. (e) UMAPs showing distribution of expression of select transcripts in the blood of Mouse 4 (Top) and Mouse 5 (Bottom). Genes include *Pdcd1* (encoding PD-1), *Klrk1* (encoding

NKG2D)*,* *Entpd1* (encoding CD39)*,* and *Cx3cr1* (encoding CX3CR1)*.* (f) Representative FACS contour plots showing all possible pairwise combinations of NKG2D, CD39, and CX3CR1 expression (as indicated in each plot) on $CD8^+$ T cells in the blood of mice at day 21 post-implantation of MC38 tumor cells. Plots are gated on singlets, live/dead$^-$, $CD45^+$, $CD8\alpha^+$. Numbers on plots indicate the percent of cells within each quadrant of the total parent population. (g) Quantification of the flow cytometry plots in (f), showing the frequency of cells expressing one, two, or three of the indicated proteins (NKG2D, CD39, and CX3CR1) determined using Boolean gating, of the population of cells expressing at least one of the markers. Shown are the average frequencies of all possible combination gates from 6 mice. In (g), 70.5% only expressed one of the markers but not the others, 20.1% expressed only two of the markers, and 9.4% expressed all three of the markers. Data are representative of three independent experiments with 5-9 mice per experiment. (h-i) Quantification of the frequencies of cells expressing one, two, or three of the indicated proteins (NKG2D, CD39, and CX3CR1) of the population of cells expressing at least one of the markers in the blood of Mouse 4 and Mouse 5 using the CITE seq data. (h) The frequencies of all possible combination gates on (h) the total population of cells from the CITE seq experiment (not subsetting based on TM status) and (i) only the TM population. In (h), 61.9% of cells expressed only one marker, 28.7% expressed only two markers, and 9.4% expressed all three markers (values averaged between Mouse 4 and 5). In (i), 28.1% of cells expressed only one marker, 52.7% expressed only two markers, and 19.2%, expressed all three markers (values averaged between Mouse 4 and 5). The pie charts in (g-h) share the legend to the left of (i).

Figure S3

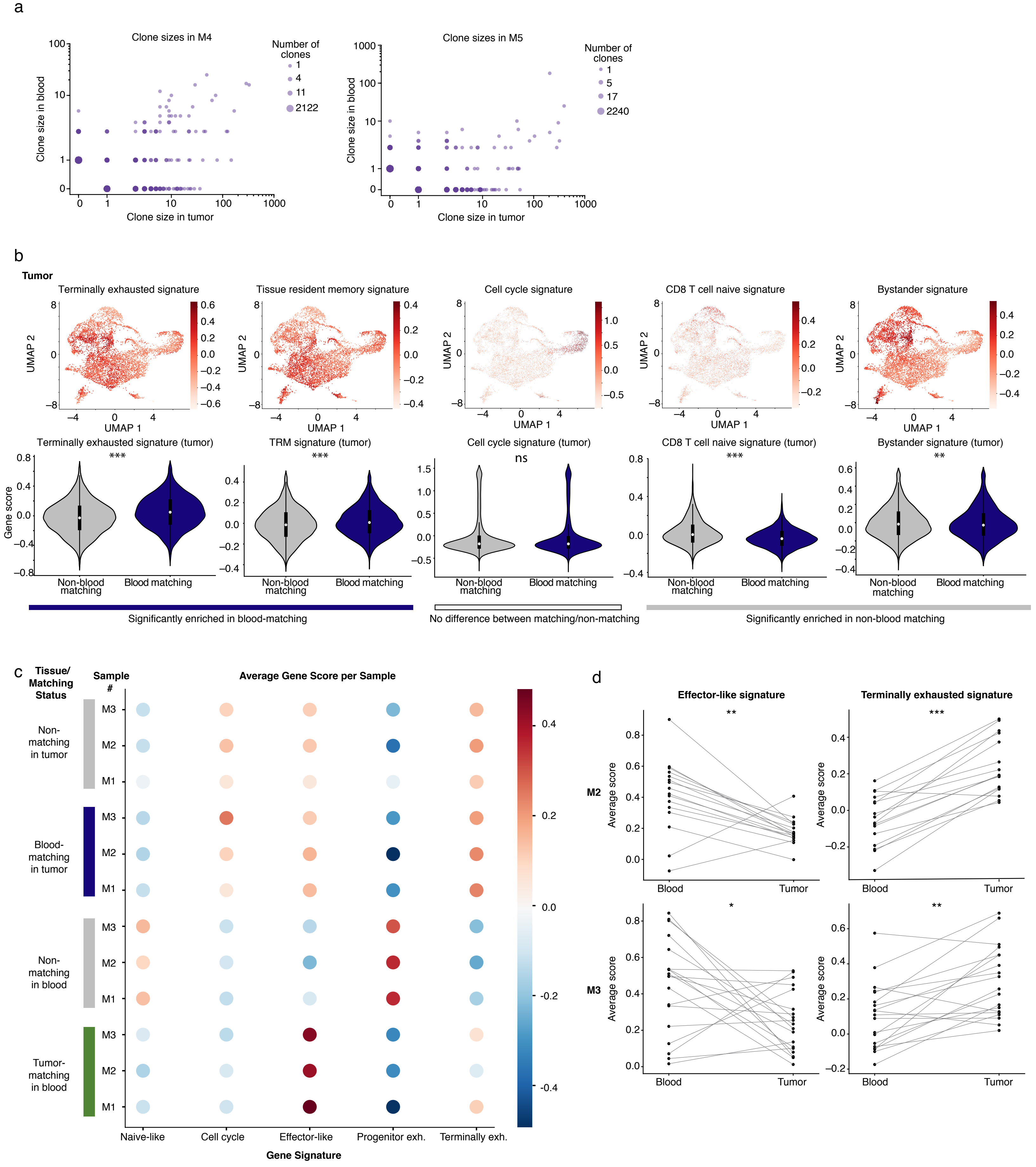
a
Clone sizes in M4
Clone sizes in M5
Number of clones
1
4
11
2122
5
17
2240
Clone size in blood
Clone size in tumor
b
Tumor
Terminally exhausted signature
Tissue resident memory signature
Cell cycle signature
CD8 T cell naive signature
Bystander signature
UMAP 1
UMAP 2
Terminally exhausted signature (tumor)
TRM signature (tumor)
Cell cycle signature (tumor)
CD8 T cell naive signature (tumor)
Bystander signature (tumor)
Gene score
***
ns
**
Non-blood matching
Blood matching
Significantly enriched in blood-matching
No difference between matching/non-matching
Significantly enriched in non-blood matching
c
Tissue/ Matching Status
Sample #
Average Gene Score per Sample
Non-matching in tumor
Blood-matching in tumor
Non-matching in blood
Tumor-matching in blood
M3
M2
M1
Naive-like
Cell cycle
Effector-like
Progenitor exh.
Terminally exh.
Gene Signature
d
Effector-like signature
Terminally exhausted signature
M2
M3
Average score
Blood
Tumor
*

**Figure S3: Tumor-matching CD8+ T cells in the blood show stronger enrichment for effector signatures and weaker enrichment for exhaustion signatures than the corresponding clones in the tumor microenvironment.** (a) Expansion rates of clones in blood and MC38 tumor (log-scale), for Mouse 4 (left) and Mouse 5 (right). (b) (Top) UMAP visualization of signatures related to CD8+ T cell transcriptional states in the mouse integrated MC38 tumor samples. From left to right are signatures of terminal exhaustion from (Miller et al., 2019), TRM cells from (Beura et al., 2018), cell cycle from (Kowalczyk et al., 2015), naïve cells from (Kaech et al., 2002), and "bystander" cells with TCRs that are not specific to the tumor from (Mognol et al., 2017). (Bottom) Violin plots quantifying the expression of each signature in blood-matching compared to non-blood matching clones. Significance determined using a Wilcoxon rank-sum test. Colored bars beneath the violin plots indicate whether the mean is statistically greater in blood-matching cells (terminal exhaustion $p=1.8x10^{-40}$, Trm $p=1.1x10^{-12}$), not statistically significant (cell cycle, p=0.97), or statistically greater in non-blood matching cells (naïve $p=2.2x10^{-65}$, bystander p=0.006). (c) Shown are average gene scores per sample for mouse blood and tumor, separated by matching status. M1-3 indicates each mouse sample number. For a given signature, a gene score was calculated for each cell. Shown are naïve-like (Kaech et al., 2002), cell cycle (Kowalczyk et al., 2015), and the effector-like, progenitor, and terminally exhausted signatures from (Miller et al., 2019). (d) Clone by clone analysis examining the mean expression of an "effector-like" gene signature or a "terminal exhaustion" gene signature from (Miller et al., 2019). Each dot shows the average gene signature of the cells in a given clone, and lines connect the same clone between blood and tumor samples. Shown are clones detected in Mouse 2 (Top) and Mouse 3 (Bottom). Significance determined using a Wilcoxon signed-rank test. For Mouse 2, $p=6.1x10^{-3}$ for the effector-like signature, and $p=5.3x10^{-4}$ for the terminally-exhausted signature. For Mouse 3, p=0.033 for the effector-like signature, and $p=1.1x10^{-3}$ for the terminally-exhausted signature.

### Figure S4

a

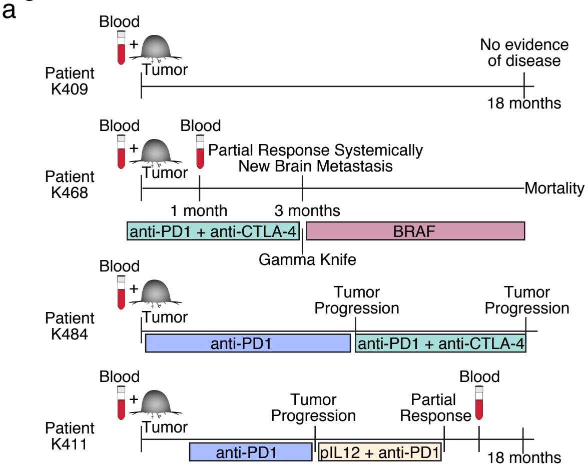


b

| Human Source | # of cells (GEX) | # of cells (GEX+TCR) | # Matching cells | % Matching cells |
|---|---|---|---|---|
| K409 blood | 5119 | 1565 | 19 | 1.21 |
| K409 LN met | 4371 | 2542 | 568 | 22.34 |
| K409 primary tumor | 1989 | 855 | 355 | 41.52 |
| K411 blood | 2543 | 218 | 60 | 27.52 |
| K411 blood longitudinal | 18524 | 4271 | 189 | 4.43 |
| K411 LN met | 3737 | 2291 | 207 | 9.04 |
| K468 blood | 8919 | 2237 | 577 | 25.79 |
| K468 blood longitudinal | 19080 | 15650 | 3107 | 19.85 |
| K468 axillary subcu. mass | 4878 | 3963 | 724 | 18.27 |
| K484 blood | 5252 | 1308 | 334 | 25.54 |
| K484 LN met | 1903 | 846 | 124 | 14.66 |

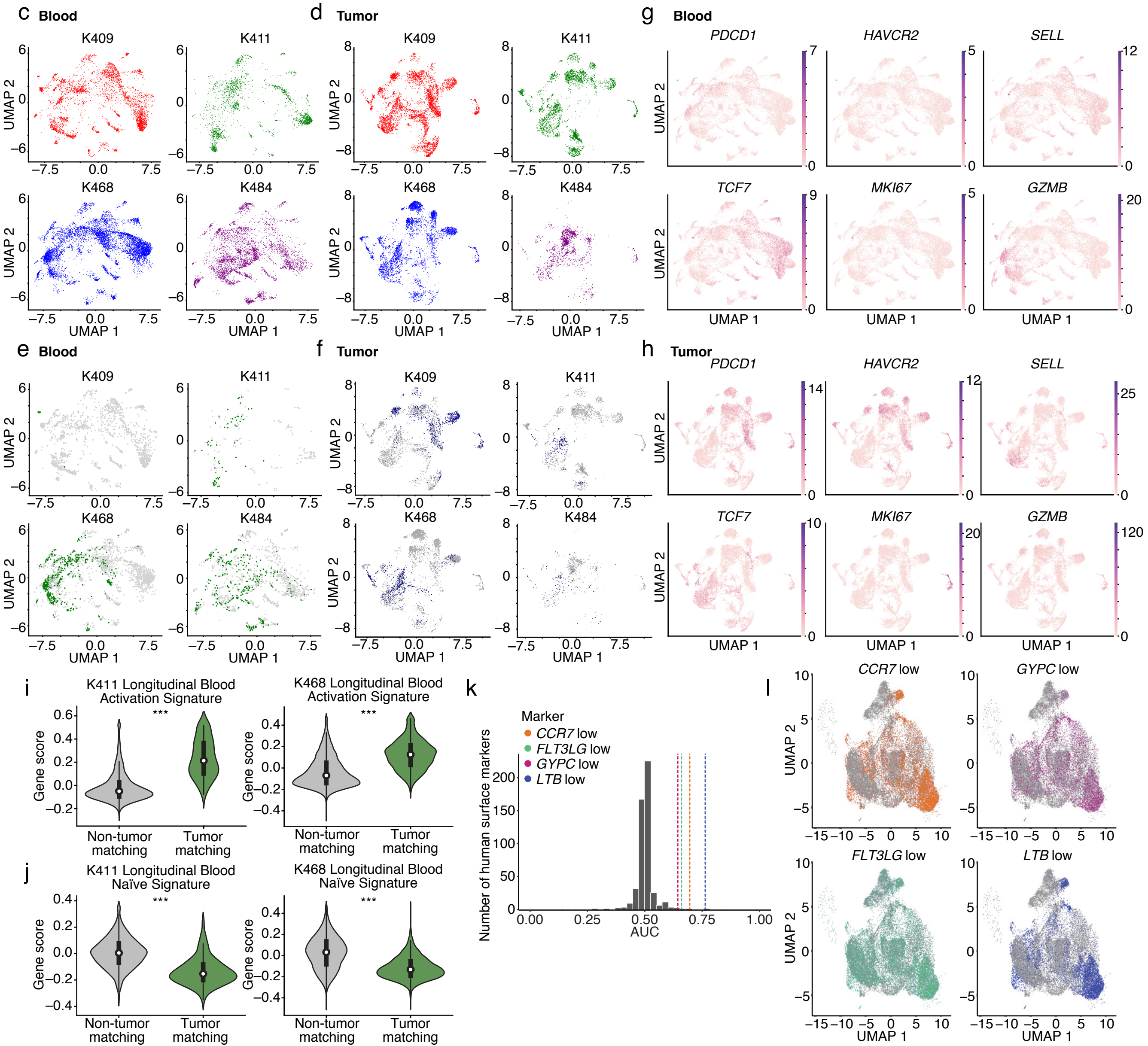

**Figure S4: Transcriptional landscape of CD8⁺ T cells in paired patient peripheral blood and melanoma samples.** (a) Schematic of clinical parameters for patient samples. Patients were checkpoint-treatment naïve at the time of initial paired blood/tumor sampling. Subsequent course of treatment indicated. Timing of longitudinal blood sample collection for follow up analysis in patients K468 and K411 indicated. The longitudinal sample for K468 was taken one month after the initial blood sample, and during that time the patient received anti-PD-1 and anti-CTLA-4 combination therapy. The longitudinal sample for K411 was taken approximately 16 months after the initial sample, after the patient had received anti-PD-1 as a single agent followed by combination therapy with anti-PD-1 and TAVO (tavokinogene telseplasmid, (Algazi et al., 2020) for TAVO monotherapy, and clinicaltrials.gov reference NCT03132675 for combination). (b) Table indicating details regarding each patient in the cohort, including the site of tissue resection, number of cells recovered that had gene expression (GEX) data, GEX and TCR data, number of matching cells, percentage matching cells of the total sorted population, and the frequency of *PDCD1*⁺ TM cells. UMAP visualization of the integrated initial paired blood samples (c and e) and melanoma samples (d and f) showing the distribution of each patient in the integrated object. Cells are colored by patient, and the remaining cells in the integrated object are excluded from visualization. (c-d) indicate all cells from a given patient, (e-f) show matching cells colored in green (TM cells in blood) or navy blue (blood-matching cells in tumor) and non-matching cells in grey from each patient. (g-h) UMAP visualizations showing the distribution of expression of select transcripts in the integrated blood (g) and melanoma (h) samples. Genes include *PDCD1* (encoding PD-1), *HAVCR2* (encoding Tim-3), *SELL* (encoding CD62L), *TCF7* (encoding TCF-1), *MKI67* (encoding Ki-67), and *GZMB* (encoding granzyme B). (i-j) Violin plots showing expression of activation (i) or naïve (j) CD8⁺ T cell signatures in TM and non-TM cells in the longitudinal blood samples from K411 and K468. Signatures derived from (Akondy et al., 2017). Significance determined used a Wilcoxon rank-sum test. For the activation signature in (i), $p=1.2x10^{-76}$ for K411 and $p<0.001$ for K468. For the naive signature in (j), $p=6.5x10^{-52}$ for K411 and $p<0.001$ for K468. (k) Histogram showing the distribution of AUC values averaged across the six patient samples for each of the human surface markers (Chihara et al., 2018). Colored lines represent the AUC for $CCR7^{low}$, $FLT3LG^{low}$, $GYPC^{low}$, and $LTB^{low}$ averaged across the six patient samples. (l) UMAP visualizations of the top singleton marker gates in human in $CD8^+$ cells from all patient blood samples integrated as described in Methods. In each plot, cells are colored if they pass the particular negation gate; that is, if they are selected as tumor-matching because their expression of the negation marker (indicating "low" expression, labeled as "$marker^{low}$"). For $CCR7^{low}$, Sensitivity = 0.827, Specificity = 0.621; for $FLT3LG^{low}$, Sensitivity = 0.780, Specificity = 0.447; for $GYPC^{low}$, Sensitivity = 0.340, Specificity = 0.819; for $LTB^{low}$, Sensitivity = 0.718, Specificity = 0.768. For patient samples, "tumor" in the Figure refers to both resections from the primary tumor and metastases as indicated in Fig. S4B.

Figure S5

a TCR cluster-based matching

| % change in number of TM cells | | | | % of blood cells added to TM compartment | | | |
|---|---|---|---|---|---|---|---|
| K409 | K411 | K468 | K484 | K409 | K411 | K468 | K484 |
| 5.26% (of 19 cells) | 6.67% (of 60 cells) | 8.15% (of 577 cells) | 12.9% (of 334 cells) | 0.06% (of 1565 cells) | 1.83% (of 218 cells) | 2.1% (of 2237 cells) | 3.29% (of 1308 cells) |

b

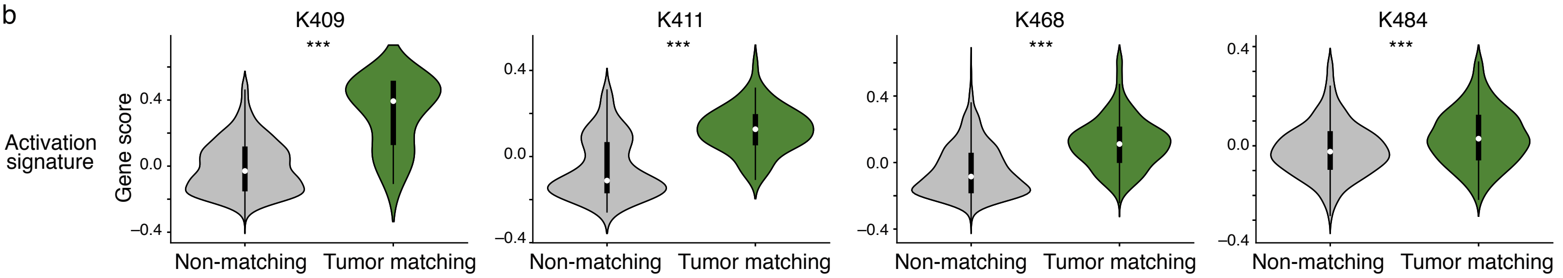


c

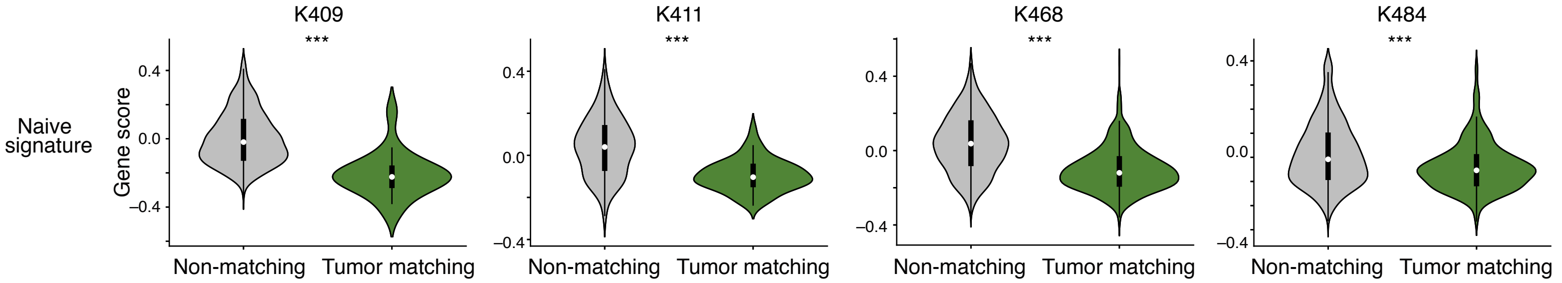


d

| | PD1 sensitivity k409 | PD1 specificity k409 | PD1 sensitivity k411 | PD1 specificity k411 | PD1 sensitivity k468 | PD1 specificity k468 | PD1 sensitivity k484 | PD1 specificity k484 | **PD1 sensitivity range** | **PD1 specificity range** |
|---|---|---|---|---|---|---|---|---|---|---|
| **exact matching** | 0.684 | 0.984 | 0.033 | 0.99 | 0.246 | 0.896 | 0.039 | 0.962 | 0.033 – 0.684 | 0.896 – 0.99 |
| **TCR cluster-based matching** | 0.722 | 0.985 | 0.047 | 1.00 | 0.244 | 0.899 | 0.043 | 0.963 | 0.043 – 0.722 | 0.899 – 1.00 |

e

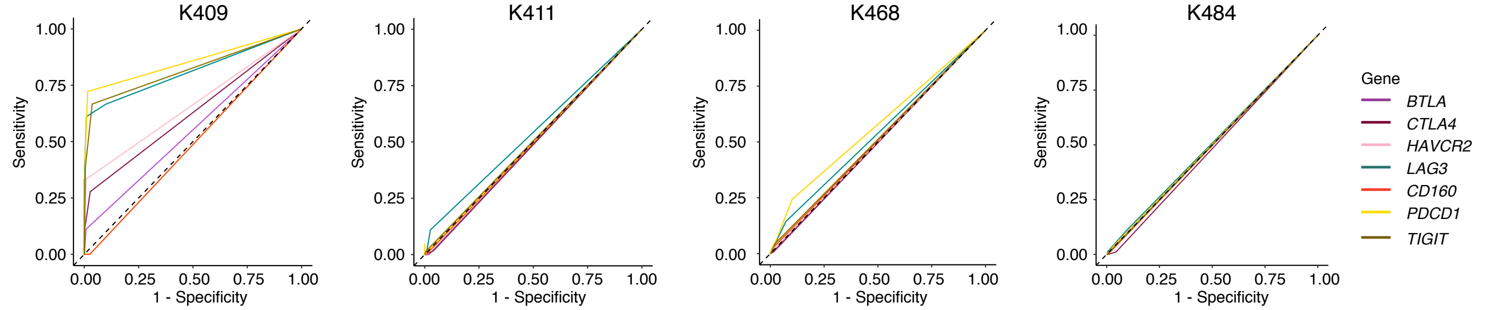


f

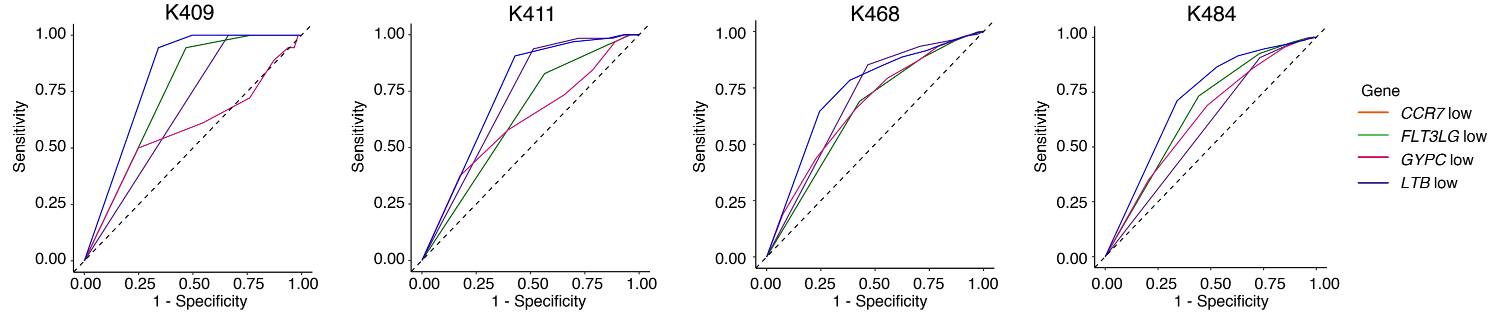

**Figure S5: Tumor-matching CD8$^{+}$ T cells identified using GLIPH2 and iSMART show similar signs of activation and sensitivity/specificity rates of inhibitory receptors and candidate surface markers as matching cells identified based on sequence matching.** (a) Summary metrics showing the increase in frequency of CD8$^{+}$ T cells classified as tumor-matching cells in each of the four primary patient samples determined using the TCR cluster-based matching method (defined as cells identified as TM using both GLIPH2 and iSMART) compared to the exact sequence matching method. (b-c) Violin plots showing enrichment of (b) activation or (c) naïve CD8$^{+}$ T cell signatures in TM and non-TM cells on the cells identified using the TCR cluster-based matching method. Signatures derived from (Akondy et al., 2017). Significance determined used a Wilcoxon rank-sum test. For the activation signature in (b), $p=3.2 \times 10^{-8}$ (K409), $p=2.6 \times 10^{-15}$ (K411), $p=8.7 \times 10^{-98}$ (K468), $p=8.5 \times 10^{-14}$ (K484). For the naive signature in (c), $p=1.5 \times 10^{-8}$ (K409), $p=3.2 \times 10^{-12}$ (K411), $p=6.1 \times 10^{-87}$ (K468), $p=3.4 \times 10^{-12}$ (K484). (d) Summary metrics showing the sensitivity and specificity of the *PDCD1* transcript to identify TM cells from non-TM cells in the blood using exact sequence matching compared to TCR cluster-based matching. (e-f) ROC curves for TM cells classified using the TCR cluster-based matching showing the sensitivity and specificity of (e) a collection of inhibitory receptor genes (*PDCD1, BTLA, CTLA4, HAVCR2, LAG3, CD160, and TIGIT*) or (f) the consensus markers for identifying TM cells (*CCR7*$^{low}$, *LTB*$^{low}$, *GYPC*$^{low}$, or *FLT3LG*$^{low}$, referred to as negation markers), shown for each patient.
